# Human IRAK kinases differentially alter metabolic regulation and mitochondrial function in *Saccharomyces cerevisiae*

**DOI:** 10.1101/2025.08.06.668917

**Authors:** Elba del Val, Giulia Genna, Teresa Fernández-Acero, María Molina, Víctor J. Cid

## Abstract

Interleukin-1 receptor-associated kinases (IRAKs) are Ser/Thr protein kinases characterized by an N-terminal Death domain (DD). Upon stimulation of Toll-like receptors (TLRs) or the interleukin-1 receptor (IL-1R), IRAKs are recruited to supramolecular signalling complexes, known as myddosomes, through interactions between their DDs and the adaptor protein MyD88. Myddosomes are essential for the activation of nuclear factor kappa B (NF-κB) in response to diverse pathogen– and damage-associated molecular patterns (PAMPs and DAMPs), and they contribute to inflammation, cell survival, and proliferation. In the hierarchical assembly of the myddosome, MyD88 first recruits IRAK4, which serves as a scaffold for the subsequent binding of IRAK1 and/or IRAK2. To explore alternative models for studying IRAK function, we expressed human IRAK1, IRAK2 and IRAK4 individually in *Saccharomyces cerevisiae* and performed a comparative analysis. Heterologous expression of these kinases, especially IRAK4, impaired yeast growth; an effect dependent on its kinase activity. Transcriptomic and biochemical assays revealed that IRAK1 and IRAK4, but not IRAK2, differentially impacted metabolic regulation. Notably, IRAK4 induced mitochondrial fragmentation and mitochondrial membrane potential depolarization, whereas IRAK1 had the opposite effect. Additionally, IRAK4 led to actin depolarization and vacuole fragmentation. Based on these findings, we develop two yeast-based bioassays to screen for IRAK4 kinase inhibitors: one based on growth recovery and another using a fluorescent reporter. We provide proof-of-concept that both assays are suitable for evaluating IRAK4 function and its pharmacological inhibition.

**Importance:** IRAK kinases are essential components of the myddosome signalling complex, a key mediator of the innate immune response, with IRAK4 playing a pivotal role in the recruitment of IRAK1 and/or IRAK2. Although the precise cellular functions of IRAK-dependent phosphorylation remain incompletely understood, IRAK inhibitors are emerging as promising therapeutic candidates for the treatment of autoimmune disorders, such as rheumatoid arthritis, and various cancers, including acute myeloid leukemia. To date, most insights into IRAK function have been derived from studies in animal models, particularly mice. Our work establishes *Saccharomyces cerevisiae* as a convenient and genetically tractable platform for *in vivo* analyses of human IRAKs. We demonstrate that IRAK kinases induce metabolic deregulation and growth inhibition in yeast. Engineered *S. cerevisiae* strains could therefore be exploited for the preclinical screening of anti-inflammatory and antitumor compounds targeting IRAK signalling, potentially contributing to the development of therapies for severe human diseases.

## INTRODUCTION

Innate immunity constitutes the first line of defense against infection and cellular damage. Innate immune cells, particularly macrophages and dendritic cells, express pattern recognition receptors (PRRs) on their surfaces, which have evolved to detect conserved pathogen– and damage-associated molecular patterns (PAMPs and DAMPs). Upon stimulation, PRRs initiate a rapid non-specific immune response. Among these receptors is the family of Toll-like receptors (TLRs), which is probably the most extensively studied. Ten TLRs (TLR1-10) have been described in humans, with TLR1, 2, 4, 5, 6, and 10 localized at the plasma membrane, and TLR3, 7, 8 and 9 at endosomal membranes(1–4).

With the exception of TLR3, all TLRs signal through the adaptor protein MyD88, which features a C-terminal Toll/interleukin-1 receptor (TIR) domain that mediates interaction with TLRs, and an N-terminal Death domain (DD) that recruits the DD-containing Interleukin-1 receptor associated kinase 4 (IRAK4). IRAK4 then acts as a scaffold for the subsequent recruitment of IRAK1 and/or IRAK2. The resulting assembly forms a supramolecular organizing center (SMOC) known as the myddosome, which functions as a dynamic platform for the assembly of additional kinases and ubiquitin ligases, ultimately leading to NF-kB nuclear translocation and activation of target gene expression (5–8).

While IRAK proteins are essential for myddosome formation, their scaffolding role is generally considered more relevant than their kinase activity. The crystal structure of a MyD88–IRAK4–IRAK2 complex revealed a helical oligomeric core composed of six MyD88, four IRAK4, and four IRAK2 subunits assembled via DD-DD interactions (9). Nevertheless, the kinase activity of IRAK4 and IRAK1 may play auxiliary regulatory roles in fine-tuning myddosome dynamics, downstream signaling, or immunometabolism (10). Dysregulated TLR signaling is associated with several autoimmune and neurodegenerative diseases, as well as with cancer syndromes, such as myeloid neoplasms (11, 12), highlighting the potential of IRAKs as drug targets. In particular, alternative splicing of IRAK4 has been observed in acute myeloid leukemia (AML), where pharmacological inhibition of IRAK4 has proven effective in reducing malignant proliferation (13). Specific IRAK4 inhibitors, such as emavusertib, are currently under clinical evaluation for the treatment of myeloid malignancies (14). Therefore, the identification of novel IRAK inhibitors holds significant therapeutic promise.

Building on our previous work with other components of the TLR signaling pathway (15, 16), we employ the single-celled eukaryotic *Saccharomyces cerevisiae* as a model system to study human IRAK proteins. Transcriptomic profiling reveals that heterologous expression of these kinases –particularly IRAK4– disrupts glucose sensing and alters multiple aspects of yeast physiology, ultimately leading to growth inhibition. Furthermore, we leverage these findings to develop yeast-based bioassays for the functional characterization and pharmacological inhibition of IRAK4 kinase activity, with relevance for the preclinical screening of compounds targeting IRAK4.

## RESULTS

### Human IRAK proteins negatively affect *S. cerevisiae* growth and viability

IRAK4, IRAK1, and IRAK2 are homologous proteins that share a conserved domain architecture, although IRAK4 lacks the C-terminal extension containing TRAF6-binding motifs (**Fig. 1A**). To investigate their effects in yeast, we cloned and expressed the corresponding human cDNAs in *S. cerevisiae* as GST fusion proteins under the control of the inducible *GAL1* promoter, which is repressed by glucose and activated by galactose. Given that IRAK4 exhibits both kinase activity-dependent and –independent functions (17), we also included in our study a catalytically inactive mutant carrying Ala substitutions at Lys residues 213 and 214 (K213A, K214A), hereafter referred to as IRAK4(KD) (for kinase-dead) (18–20).

**Figure 1.**
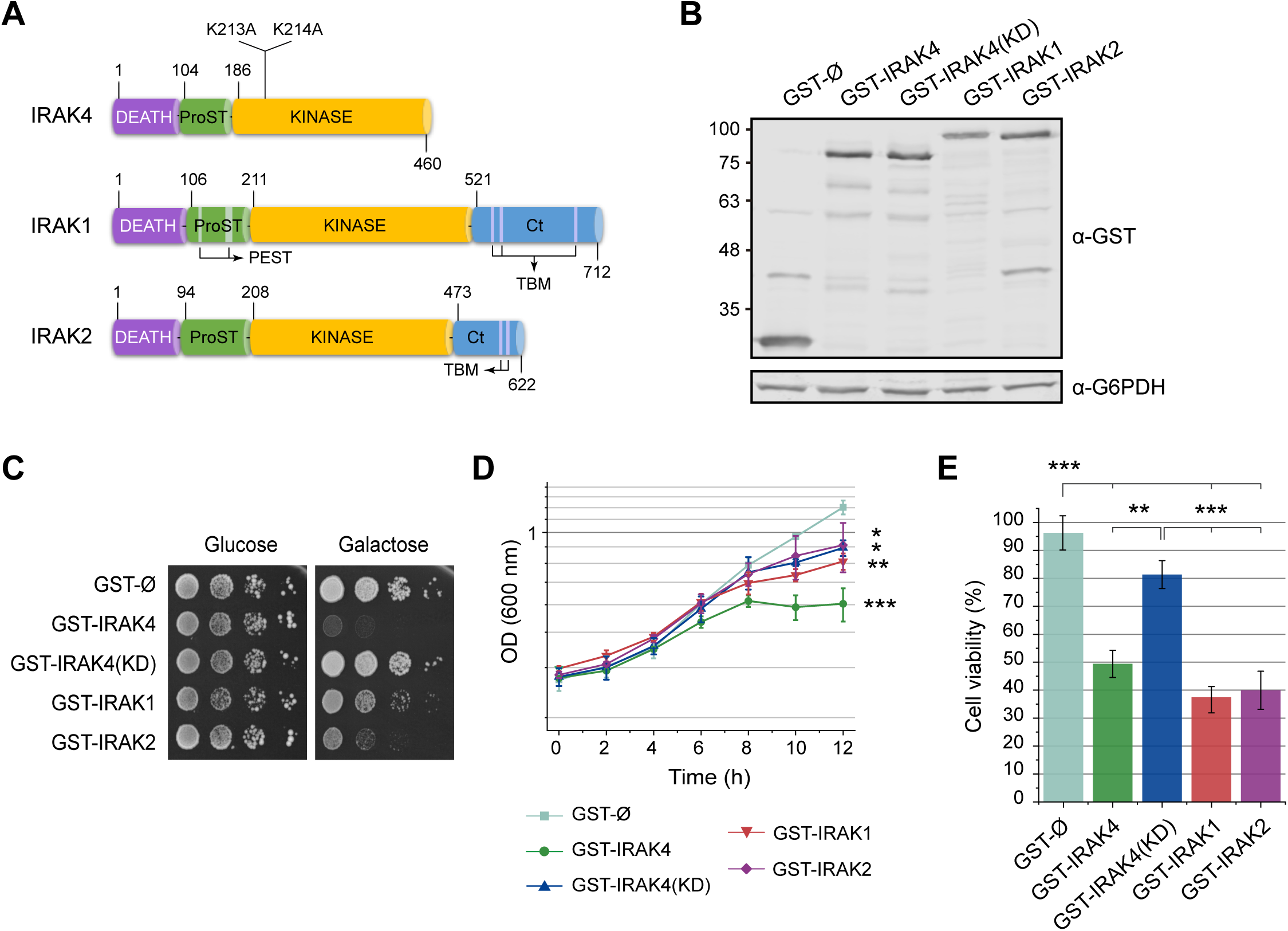
Expression of human IRAK proteins in *S. cerevisiae.* **A.** Schematic representation of IRAK domain structures. All IRAK proteins contain a Death domain, a ProST domain [rich in proline (Pro), serine (S), and threonine (T)], and a kinase domain. The C-terminal domain, which contains up to three TRAF6-binding motifs (TBM), is present in IRAK1 and IRAK2 but absent in IRAK4. Additionally, IRAK1 has two PEST sequences [rich in proline (P), glutamate (E) or aspartic acid (D), serine (S), and threonine (T)] within the ProST region. Domain boundaries are indicated above and below each diagram. **B.** Immunoblot of cell extracts from the YPH499 strain transformed with the plasmids encoding GST alone or GST fusions of IRAK4, IRAK4(KD), IRAK1, and IRAK2 under the *GAL1* promoter. Expected molecular weights are 26, 78, 78, 93, and 85 kDa, respectively. Proteins were detected using an anti-GST antibody. G6PDH was used as a loading control. **C.** Spot growth assay showing yeast growth on solid selective medium (Ura⁻ Leu⁻) supplemented with glucose (expression repressed) or galactose (expression induced). The YPH499 strain was transformed with the same plasmids as in B. Cultures were serially diluted and spotted onto plates. Results shown are representative of three biological replicates. **D.** Growth curves in liquid SG Ura⁻ Leu⁻ medium following induction for 12 h. Optical density at 600 nm (OD₆₀₀) was measured every 2 h. Data represent the mean ± SD from three biological replicates. Statistical significance at the final time point (12 h) compared to the empty vector control is indicated: p < 0.05 (*), p < 0.01 (**), and p < 0.001 (***), based on Tukey’s HSD test. **E.** Viability assay following 12 h induction in SG Ura⁻ Leu⁻ medium. Cells were plated onto YPD agar and incubated for 18 h. Colony-forming units (CFU) were counted, and viability was expressed as a percentage relative to the control strain carrying the empty vector (set to 100%). Data represent the mean ± SD of three biological replicates. Statistical significance is indicated as in panel D.

All three proteins were stably expressed when yeast clones were grown in galactose-containing medium, as verified by immunoblotting (**Fig. 1B**). However, IRAK4 expression caused a marked growth arrest in both solid (**Fig. 1C**) and liquid (**Fig. 1D**) cultures, while IRAK1 and IRAK2 led to milder but still significant growth inhibition. The toxic effect of IRAK4 in yeast was dependent on its kinase activity, as expression of IRAK4(KD) abolished toxicity on solid medium and only mildly affected growth in liquid culture, to a level comparable that of IRAK2 (**Fig. 1C-D**). To further evaluate cell viability, we assessed the ability of pre-induced clones (12 h in galactose) to form microcolonies upon transfer to glucose-containing medium. We found that IRAK expression resulted in irreversible growth arrest in approximately 50% of the cells. Furthermore, IRAK4-induced loss of viability was largely dependent on its kinase activity (**Fig. 1E**).

To determine whether the observed growth arrest and loss of viability in IRAK-expressing cells were due to cell lysis or cell cycle arrest, we performed flow cytometry analyses. A small but statistically significant percentage of yeast cells expressing active IRAK4 were propidium iodide (PI) positive (11.43 ± 0.12 %), indicating compromised membrane integrity compared to the catalytically inactive mutant. In contrast, cells expressing IRAK1, IRAK2, or IRAK4(KD) showed PI uptake levels comparable to those of the empty vector control (**Suppl. Fig. S1A**), suggesting that loss of membrane permeability is not the primary cause of both growth inhibition and reduced viability.

To assess potential cell cycle effects, we measured DNA content by flow cytometry. No significant changes in the n/2n DNA ratio were observed in any condition (**Suppl. Fig. S1B**), ruling out cell cycle arrest as the underlying cause of growth impairment. However, cells expressing active IRAK4 showed a small but significant percentage of cells with a DNA content higher than 2n, consistent with subtle defects in nuclear segregation or cytokinesis. To further explore this possibility, we examined the actin cytoskeleton using rhodamine-phalloidin staining. Budding cells expressing catalytically active IRAK4 displayed a highly depolarized actin cytoskeleton, a phenotype not observed in cells expressing IRAK4(KD), IRAK1, or IRAK2 (**Suppl. Fig. S2**). These cytoskeletal abnormalities may contribute to the pronounced growth defect associated to IRAK4 expression.

In summary, human IRAK4, IRAK1, and IRAK2 differentially impair yeast growth, with IRAK4 exerting the strongest effects through a kinase-dependent mechanism that may involve defects in actin organization. These findings support the use of *S. cerevisiae* as a platform to functionally assess IRAK activity and inhibition.

### Human IRAK kinases trigger distinct patterns of metabolic stress in yeast

To understand the nature of the cellular damage caused by human IRAK kinases in yeast, we performed transcriptomic profilling using DNA microarrays. We compared global gene expression in *S. cerevisiae* cells overexpressing GST-tagged IRAK4, IRAK4(KD), IRAK1, and IRAK2 to that in control cells expressing GST alone. All cells were harvested 6 h after switch to galactose-based medium to induce protein expression. The complete dataset is available in the GEO Omnibus repository (accession number GSE236671). Transcriptional changes measured with DNA microarrays were validated by quantitative real-time PCR (RT-qPCR) (**Suppl. Fig. S3**).

The extent of transcriptional deregulation varied greatly among IRAK proteins. Differentially expressed genes were defined as those showing average fold-changes >2 (upregulated) or <0.5 (downregulated), with a false discovery rate (FDR) <0.05 across three biological replicates (**Suppl. Tables S1-S4**). IRAK1 had the most pronounced effect, upregulating 388 genes and downregulating 150 genes (**Fig. 2A**. **Suppl. Table S1**), while IRAK2 barely altered the transcriptome (**Suppl. Table S2**). IRAK4 also led to remarkable changes in gene expression, with 180 genes upregulated and 115 downregulated (**Suppl. Table S3**). Interestingly, IRAK4(KD) also triggered modest but statistically significant gene expression changes (**Suppl. Table S4**), suggesting kinase-independent regulatory effects that were even more pronounced than those caused by IRAK2. Principal component analyses (PCA) revealed clear clustering of samples by IRAK type, with IRAK1, IRAK4, and IRAK4(KD) samples separating distinctly from one another and from the control, whereas IRAK2 samples clustered with the control (**Fig. 2B**).

**Figure 2.**
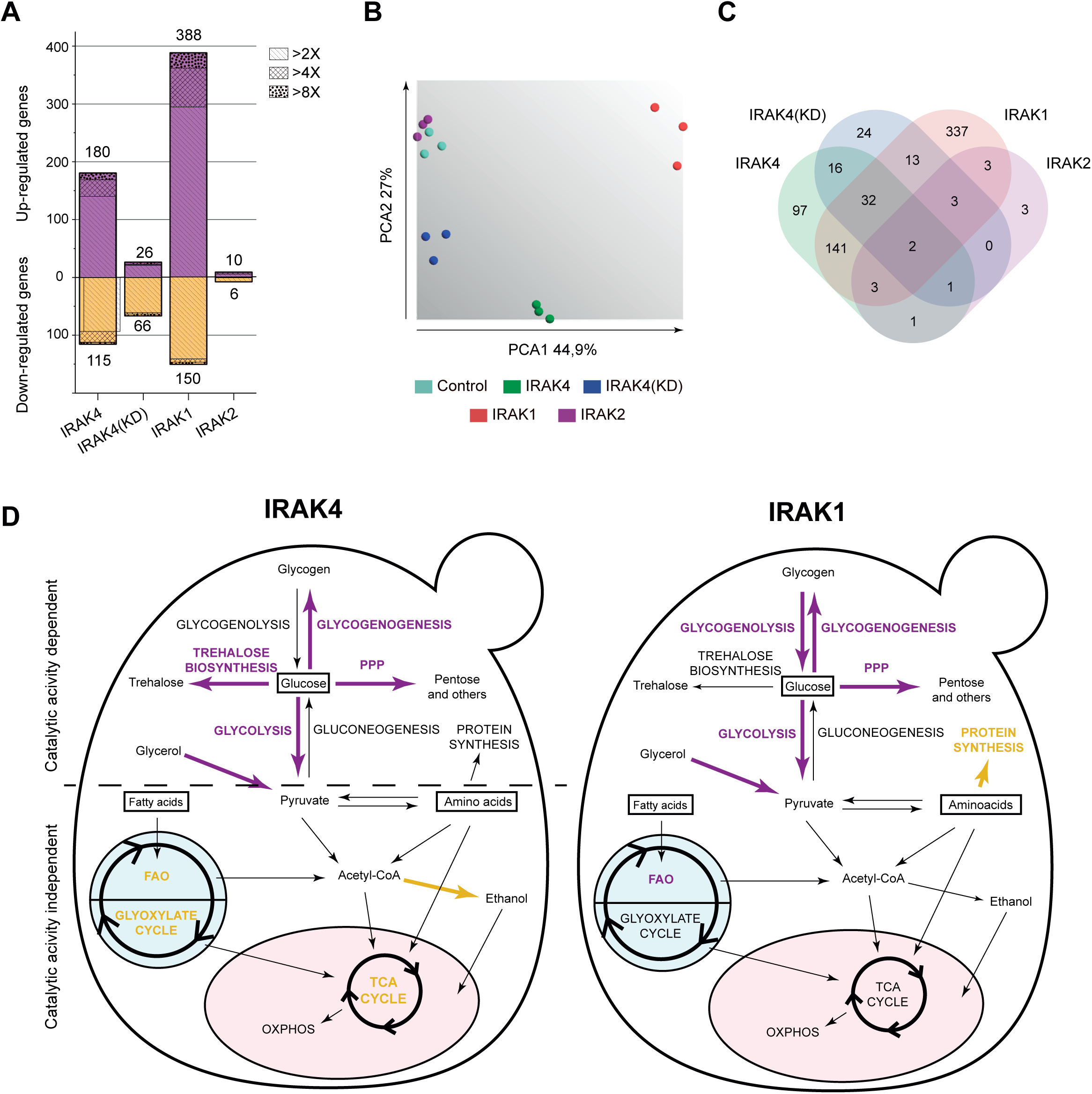
Effects of IRAK expression on the yeast transcriptome. Yeast strain YPH499 was transformed with plasmids encoding GST alone (GST-Ø, control) or GST-tagged IRAK4, IRAK4(KD), IRAK1, and IRAK2, and cultured for 6 h in SG Ura⁻ Leu⁻ medium prior to transcriptomic analysis. **A.** Bar graph showing the number of differentially expressed *S. cerevisiae* genes in each experimental condition relative to the control. Upregulated genes (FC ≥ 2) are shown in the purple above the x-axis; downregulated genes (FC ≤ 0.5) are shown in the yellow below. Bar textures indicate fold changes threshold of ≥2, ≥4, or ≥8. Only genes with a false discovery rate (FDR) <0.05 were included. **B.** Principal Component Analysis (PCA) of gene expression data across all conditions. The first two principal components (PC1 and PC2) account for 71.9% of the total variance. The plot was generated using TAC software v.4.0.2, based on the R function ‘prcomp’. **C.** Venn diagram illustrating the overlap of differentially expressed among IRAK4, IRAK4(KD), IRAK1, and IRAK2 conditions. Overlapping regions represent genes commonly deregulated in multiple conditions. **D.** Schematic representation of metabolic pathways affected by IRAK expression. Diagrams depict a yeast cell, with mitochondria (pink ellipse) and peroxisomes (blue circle). The left panel shows metabolic pathways influenced by IRAK4 and IRAK4(KD); the latter highlights pathways also repressed by wild-type IRAK4. The right panel displays pathways affected by IRAK1 expression. Transcriptionally upregulated pathways are indicated in purple; downregulated pathways are indicated in yellow.

We next examined the overlap between gene sets deregulated by IRAK1 and IRAK4. Of the genes affected by IRAK4, 48.8% were also altered by IRAK1. Conversely, 62.64% of IRAK1-regulated genes and 32.88% of IRAK4-regulated genes were unique to each dataset, respectively (**Fig. 2C**). We mined gene ontology functional annotations in the up– and downregulated gene sets. In the case of IRAK4, upregulated genes were significantly enriched for functions related to trehalose biosynthesis, carbohydrate catabolism, energy production, and response to extracellular stimuli (**Suppl. Table S5; Fig. 2D**, left). On the other hand, down-regulated genes were significantly enriched for fatty acid β-oxidation, as well as related biological processes, such as propionate metabolism and long-chain fatty acid transport. Notably, fatty acid β-oxidation was also overrepresented among downregulated genes in the IRAK4(KD) dataset (**Suppl. Table S6** and **Fig. 2D**), indicating that this repression is independent of IRAK4 catalytic activity.

In contrast, IRAK1 upregulated gene sets were enriched for fatty acid β-oxidation, trehalose and glycogen biosynthesis, late nucleophagy, and oxidative stress response. Downregulated genes were associated with anabolic processes such as amino acid biosynthesis, translation, and purine compound biosynthesis (**Suppl. Table S7** and **Fig. 2D**, right). No significant enrichment of biological processes was detected in the IRAK2 dataset. Analysis of transcription factors enrichment among dysregulated genes in the datasets revealed common regulators involved in metabolism (e.g., Yap6, Spt23, and Oaf1), and stress responses (e.g., Hot1, Sko1, Cad8, and Cad1, a paralog of Yap1) (**Suppl. Table S8**).

Finally, we compared our data with existing transcriptional datasets related to metabolic and environmental stress in yeast (21) (**Suppl. Fig. S4**). The expression profile induced by IRAK4 showed strong similarity to that of cells exposed to heat stress, amino acid starvation, or short-term (1 h) nitrogen depletion. In contrast, IRAK1 expression resembled transcriptional responses to long-term (2 days) nitrogen starvation, oxidative stress (e.g., diamide or hydrogen peroxide), and the diauxic shift (**Suppl. Fig. S4**). In sum, these results indicate that IRAK kinases elicit metabolic stress-like transcriptional programs in yeast, with IRAK1 and IRAK4 triggering distinct adaptive responses.

### IRAK4 alters metabolic regulation in yeast

Most transcriptomic changes observed in cells expressing IRAK4 or IRAK1 were related to metabolic regulation, particularly glycogen and trehalose biosynthesis. These energy storage polymers typically accumulate in yeast cells under starvation or nutrient-limiting conditions (22). Since the cells in our experiments were not starved, we hypothesized that IRAK expression may impair nutrient sensing, triggering a starvation-like transcriptional response despite the presence of excess carbon and nutrients. To test this, we assessed glycogen accumulation in IRAK-expressing cells using iodine staining. As shown in **Fig. 3A**, cells expressing IRAK4 and IRAK1 —but not IRAK4(KD)— accumulated glycogen to levels comparable to those of starved cells, under non-starving conditions. IRAK2 was excluded from this analysis due to its negligible transcriptional impact.

**Figure 3.**
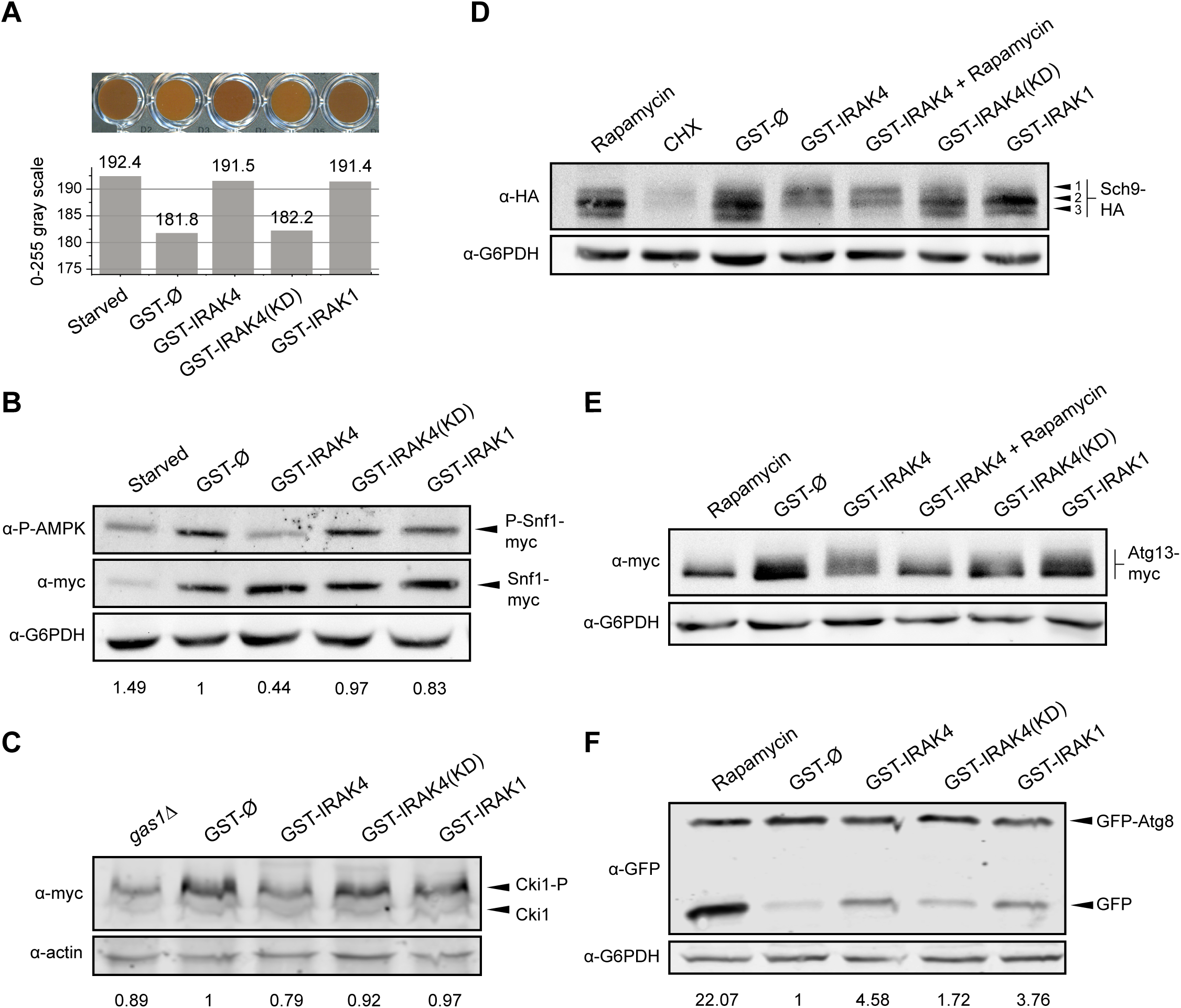
IRAK4 expression induces glycogen accumulation, deregulates Snf1 and TORC1 signalling, and promotes autophagy in yeast. **A.** Glycogen accumulation assay in *S. cerevisiae* strain YPH499 transformed with plasmids encoding GST alone (GST-Ø), GST-IRAK4, GST-IRAK4(KD) or GST-IRAK1. Cells were cultured for 8 h in SG Ura^−^ Leu^−^ culture medium. A stationary-phase culture (48 h) of the same strain served as a positive control for glycogen accumulation. **B.** Snf1 activation in strain EVY7 (Snf1-Myc) transformed with the same plasmids and incubated under the same conditions as in panel A. Cells subjected to carbon starvation (synthetic medium without sugar, 6 h) served as positive controls for AMPK/Snf1 activation. Total Snf1-Myc was detected with anti-Myc antibody; phosphorylated Snf1 was detected using a phospho-specific anti-AMPKα (Thr172) antibody. G6PDH was used as a loading control. **C.** PKA activity was assessed in YPH499 cells co-transformed with plasmids pRS423-prCUP-6xMYC-Cki12-200(S125/130A) and the indicated GST-fusion constructs. Cells were cultured for 6 h in SG Ura^−^ Leu^−^ His^−^ medium. A *gas1Δ* mutant carrying GST-Ø was used as a PKA inhibition control. PKA activity was estimated from the ratio of phosphorylated (Cki1-P) to unphosphorylated Cki1, normalized to GST-Ø. 6xMYC-Cki1 was detected with anti-Myc; actin was used as a loading control. IRAK proteins were detected with anti-GST (not shown). **D.** TORC1 activity analysis in YPH499 cells co-transformed with pRS413-Sch9-HA and the indicated GST-fusion constructs, cultured for 8 h in SG Ura^−^ Leu^−^ His^−^ culture medium. TORC1 inhibition and activation controls were prepared by treating cells with rapamycin (250 μg/mL) or cycloheximide (CHX, 12.5 μg/mL) for the final 6 h. Sch9-HA was detected using anti-HA antibody; G6PDH was used as a loading control. **E.** Phosphorylation status of Atg13-Myc in strain EVY19 transformed with the indicated GST-fusion constructs and incubated for 8 h in SG Ura^−^ Leu^−^ culture medium. Cells treated with rapamycin (250 µg/mL, 6 h) served as TORC1 inhibition controls. Atg13-Myc was detected using anti-Myc; G6PDH was used as a loading control. **F.** Autophagic flux was assessed in YPH499 cells transformed with pRS314-GFP-Atg8 and the indicated GST-fusion constructs, cultured for 8 h in SG Ura^−^ Leu^−^ Trp^−^ culture medium. Rapamycin-treated cells (100 nM, 6 h) were used as a positive control for autophagy induction. GFP-Atg8 protein and released GFP were detected using anti-GFP antibody. G6PDH was used as a loading control. Quantification represents the ratio of free GFP signal to G6PDH, normalized to the GST-Ø condition. All experiments were performed in biological triplicates; representative results are shown. In A, B and C, electrophoresis gels were prepared with 8% acrylamide-bisacrylamide and 1.5 mm thickness; gels were run at 50 V for the first 30 min, followed by 100 V.

Snf1, the yeast homolog of mammalian AMP-activated protein kinase (AMPK), is a central regulator of carbon metabolism. Under low glucose conditions, it becomes phosphorylated and promotes gluconeogenesis, respiratory metabolism, glycogen accumulation, and autophagy, while repressing anabolic processes such as fatty acid and amino acid biosynthesis (23). Several of these metabolic features were reflected in the transcriptomic profiles of IRAK1– and IRAK4-expressing cells. Since galactose was used as the carbon source to induce expression from the *GAL1* promoter, Snf1 would be expected to be active under these conditions to support the galactose switch and glucose repression (24). We measured Snf1 phosphorylation using phospho-specific anti-AMPK antibodies and compared log-phase cultures of IRAK-expressing cells to both the empty vector control and starved cells. Total Snf1 protein levels were reduced under starvation, while in IRAK-expressing log-phase cultures in galactose they remained comparable to those of the control. As expected, in starving cells, Snf1 was phosphorylated. However, phosphorylated Snf1 was diminished in IRAK4-expressing cells, but not in those expressing IRAK4(KD) or IRAK1 (**Fig. 3B**), suggesting that IRAK4 activity interferes with Snf1 signaling.

We next examined the cAMP-PKA pathway, another key regulator of nutrient signaling and metabolic control, which negatively regulates glycogen accumulation (25). Unlike Snf1, PKA activity is expected to remain high during log-phase growth in both glucose and galactose media (26). To assess PKA activity, we used PKA-specific phosphorylation of the choline kinase Cki1 as a readout (27). IRAK4-expressing cells showed reduced phospho-Cki1 levels, comparable to those in a *gas1*Δ strain known for impaired PKA activity (28). In contrast, neither IRAK4(KD) nor IRAK1 expression had a significant effect (**Fig. 3C**).

The TORC1 pathway is a third major regulator of nutrient responses, controlling cell growth and autophagy, and functionally interacting with both Snf1 and PKA pathways (29). Moreover, there is an interplay between Snf1 activity and TORC1 activation under glucose starvation (29). We first assessed TORC1 activity by examining the phosphorylation state of Sch9, a direct TORC1 substrate (30). IRAK4 —but not IRAK4(KD) or IRAK1— induced hyperphosphorylation of Sch9, detectable as reduced mobility in SDS-PAGE, resembling the effect of cycloheximide. This shift was insensitive to rapamycin, suggesting that IRAK4 may promote Sch9 phosphorylation through a TORC1-independent mechanism (**Fig. 3D**). To further evaluate TORC1 activity, we monitored the phosphorylation state of Atg13-myc, another TORC1 substrate whose phosphorylation inhibits autophagy (31). IRAK4 expression led to a slower-migrating Atg13-myc band, indicative of hyperphosphorylation, which reverted to the faster-migrating form upon rapamycin treatment, confirming TORC1 dependence (**Fig. 3E**).

Finally, we assessed autophagic activity using the Atg8-GFP processing assay (32). Contrary to expectations based on TORC1 activation, both IRAK4– and IRAK1-expressing cells showed increased levels of free GFP, reflecting enhanced autophagic flux. However, this increase was modest compared to rapamycin-treated cells (**Fig. 3F**).

In sum, IRAK4 and IRAK1 expression in yeast led to glycogen accumulation and autophagy induction under non-starving conditions. However, IRAK4 uniquely impaired the activity of Snf1 and PKA signaling pathways and simultaneously induced TORC1 signaling, resulting in Sch9 and Atg13 hyperphosphorylation. Despite TORC1 activation, autophagy remained upregulated, suggesting a decoupling of TORC1-mediated repression. These metabolic phenotypes are consistent with a broad dysregulation of nutrient-sensing pathways, in line with our transcriptomic findings.

### Human IRAK kinases differentially affect mitochondrial morphology and function in yeast

Given the central role of mitochondria in cellular metabolism (33), we examined mitochondrial morphology in yeast cells expressing IRAK4 and IRAK1. Cells were co-transformed with plasmids encoding human IRAK proteins and the mitochondrial marker Ilv6-mCherry. Mitochondria with a continuous tubular structure were classified as normal, a phenotype predominantly observed in cells carrying the empty vector or expressing IRAK4(KD) (**Figs. 4A-B**). In contrast, this phenotype was significantly reduced in cells expressing IRAK4 or IRAK1. Specifically, 38.5 ± 8.71 % of IRAK4-expressing cells exhibited hyperfragmented mitochondria, while 55.54 ± 7.43 % of IRAK1-expressing cells displayed a branched tubular morphology (**Fig. 4B**). These results suggest that heterologous expression of IRAK kinases interfere with the fission-fusion balance that governs mitochondrial morphogenesis.

**Figure 4.**
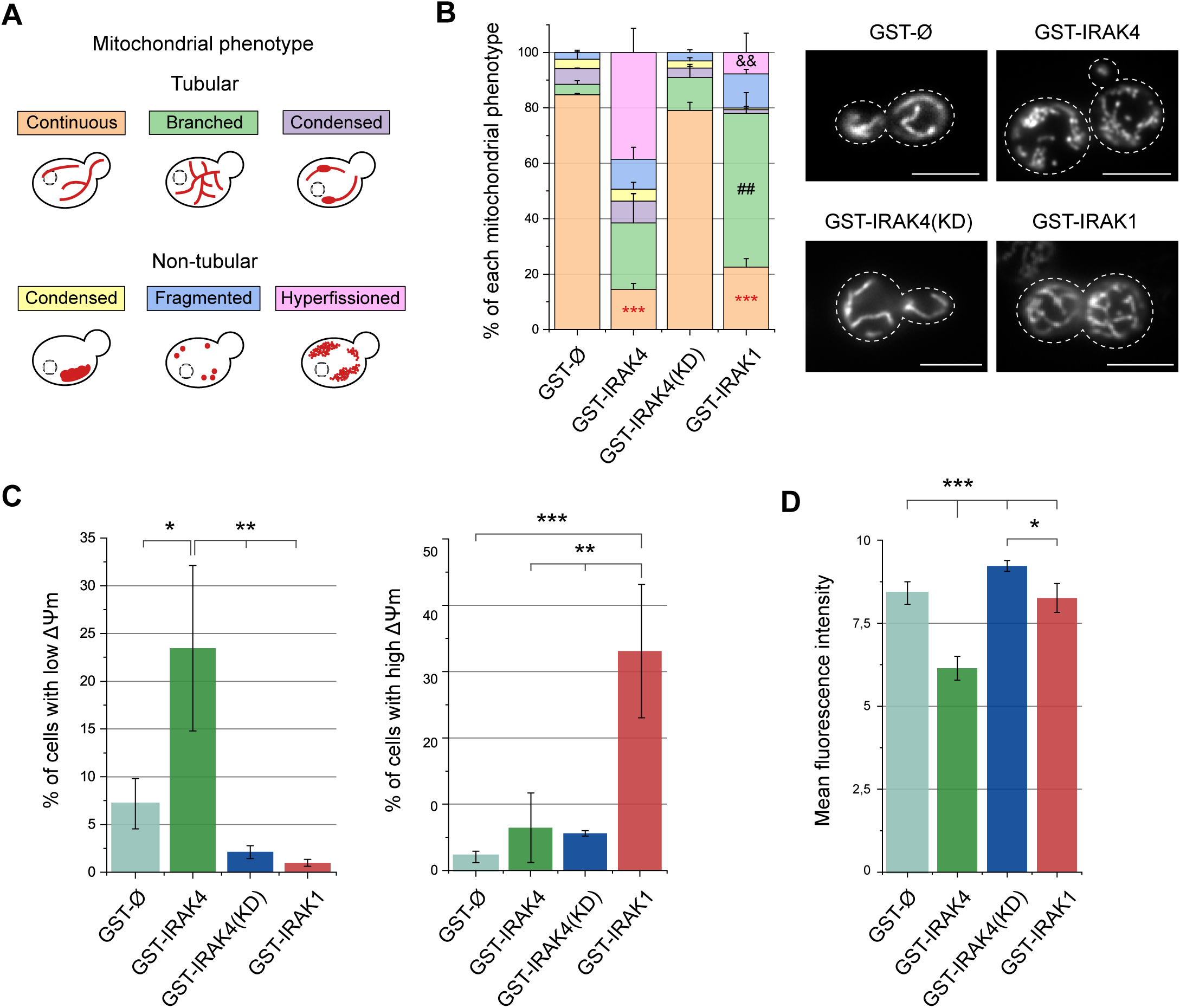
IRAK4 and IRAK1 expression alters mitochondrial morphology in *S. cerevisiae*. **A.** Schematic representation of the six mitochondrial phenotypes observed *in S. cerevisiae* following heterologous expression of human IRAK kinases or empty vector. Mitochondria (red) are shown within the outline of a yeast cell; dashed circles indicate nuclei. **B.** Left panel: stacked column graph showing the distribution of mitochondrial phenotypes in strain YPH499 co-transformed with plasmids YEplac112-Ilv6-mCherry and pEG(KG)-GST-Ø, – IRAK4, –IRAK4(KD), or –IRAK1. Cultures were grown for 12 h in SG Ura^−^ Leu^−^ Trp^−^ medium. Colors correspond to phenotypes depicted in panel A. Error bars represent standard deviation (SD). Asterisks (***) indicate significant differences in the prevalence of the continuous tubular phenotype between the control and cells expressing IRAK4 or IRAK1 (p <0.001, according to Tukey’s HSD test). Hash marks (##) and ampersands (&&) denote significant differences between IRAK4 and IRAK1 for the branched tubular and hyperfissioned non-tubular phenotypes, respectively (p < 0.01, Tukey’s HSD test). The experiment was performed in triplicate, with >100 cells analyzed per replicate. Right panel: representative fluorescence microscopy images for each condition. Scale bar: 5 μm. **C.** Quantification of mitochondrial membrane potential (ΔΨm) by rhodamine 123 (Rd123) staining and flow cytometry. Bar graph shows the percentage of cells above or below the threshold set in the overlaid histogram, representing hyperpolarized and depolarized mitochondria, respectively. Error bars show SD. Statistical significance is indicated as p < 0.05 (*), p < 0.01 (**), and p < 0.001 (***), according to Tukey’s HSD test. A representative histogram is shown in **Suppl. Fig. S5A**. **D.** Measurement of mitochondrial oxidative potential using DHR123 staining and flow cytometry. Bar graph shows the arithmetic mean of median fluorescence intensity from three biological triplicates. Error bars represent SD. Asterisks (***) and (*) indicate statistical significance at p <0.001 and p <0.05, respectively, according to Tukey’s HSD test. A representative histogram is shown in **Suppl. Fig. S5B**.

Mitochondrial membrane potential (ΔΨm) is a key indicator of mitochondrial function. To assess ΔΨm, we stained cells with rhodamine 123 (Rd 123). IRAK4 expression led to a statistically significant increase in the proportion of cells with reduced ΔΨm (i.e., mitochondrial depolarization), whereas IRAK1 expression was associated with an increased percentage of cells with elevated ΔΨm (i.e., hyperpolarization) (**Fig. 4C**). No significant differences were observed between IRAK4(KD)-expressing cells and the empty vector control. To further evaluate mitochondrial functionality, we assessed oxidative capacity using dihydrorhodamine 123 (DHR 123). IRAK4-expressing cells showed reduced ROS-dependent fluorescence compared to controls, indicating deficient mitochondrial oxidative activity (**Fig. 4D**). In contrast, no significant changes in ROS production were detected in cells expressing IRAK4(KD) or IRAK1.

Altogether, these findings indicate that IRAK4 expression impairs mitochondrial function in yeast, as evidenced by mitochondrial hyperfragmentation, membrane depolarization, and reduced oxidative capacity —effects that depend on its catalytic activity. On the other hand, IRAK1-expressing cells exhibited a distinct phenotype characterized by mitochondrial branching without loss of ROS production, suggesting that their mitochondria remain functionally competent for ATP production. Mitochondrial functional integrity may underlie the lower cytotoxicity observed for IRAK1 compared to IRAK4 in yeast cells.

### IRAK4 induces severe vacuolar fragmentation in yeast

Yeast vacuoles, functionally analogous to lysosomes in higher eukaryotes, are dynamic organelles that adapt to metabolic stages and environmental cues. To investigate the potential impact of IRAK kinases on the endocytic pathway, we used the fluorescent dye FM4-64, which is internalized via endocytosis and delivered to the vacuolar membrane in *S. cerevisiae* (34). No delays in FM4-64 uptake were observed in any of the strains, indicating that the endocytic trafficking remained functional (data not shown).

However, a distinct phenotype was observed specifically in cells expressing catalytically active IRAK4: a marked alteration in vacuolar morphology. Based on the established vacuolar protein sorting (*vps*) mutant classification (35), IRAK4-expressing cells exhibited a characteristic class B phenotype —accumulation of multiple small vacuoles— indicative of vacuolar fragmentation (**Fig. 5**). In contrast, cells expressing IRAK4(KD) or IRAK1 displayed a wild-type class A phenotype with typical vacuolar morphology. Additionally, although less frequently, IRAK4-expressing cells also showed a modest increase in phenotypes corresponding to class D (single enlarged vacuole) and class F (fragmented vacuolar structures surrounding a central vacuole), suggesting a broad disruption of vacuolar homeostasis linked to IRAK4 kinase activity.

**Figure 5.**
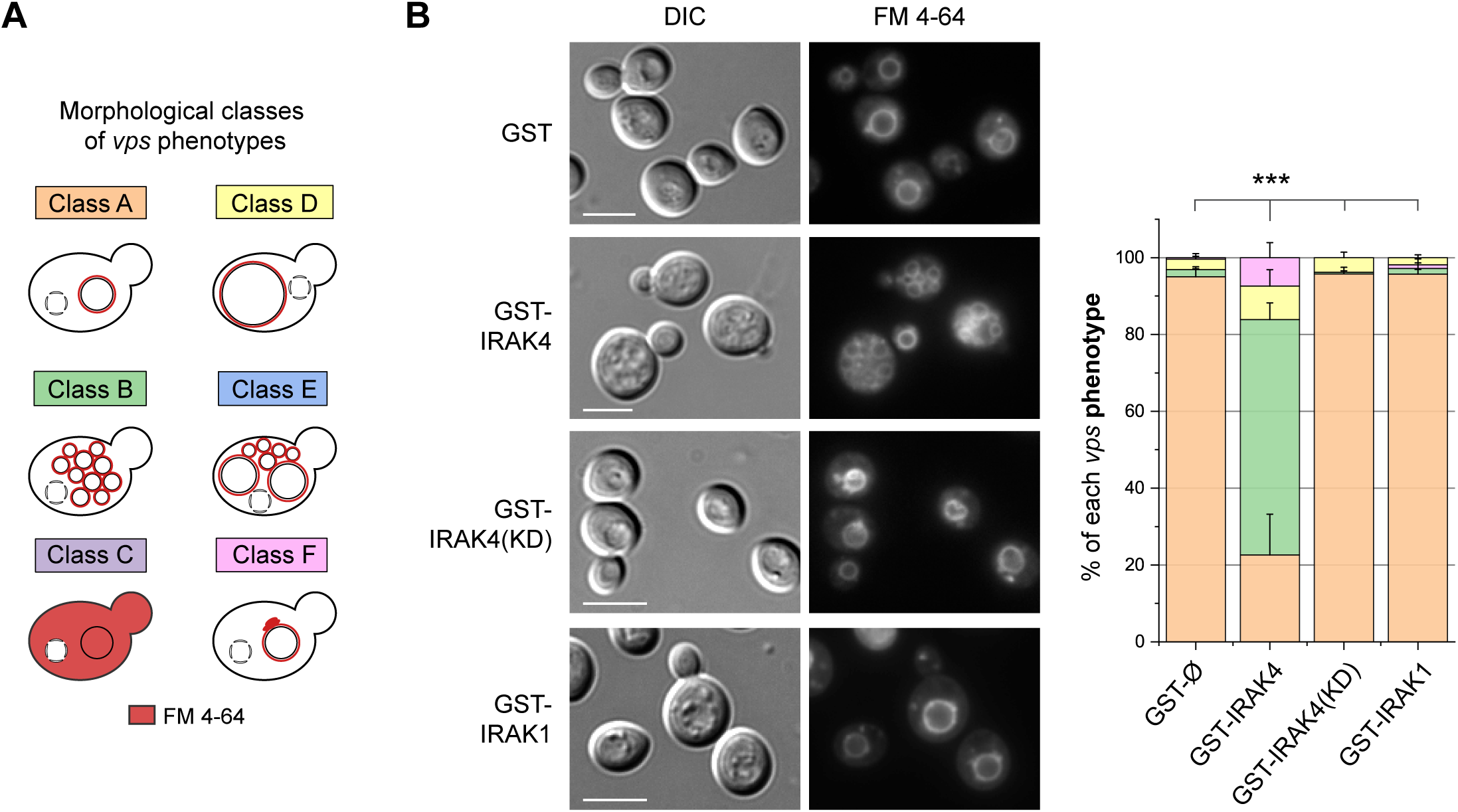
IRAK4 expression in yeast induces vacuolar fission. **A.** Schematic representation of the vacuolar morphology observed in *vps* mutant classes A-F stained with FM 4-64, based on the classification by Vida and Emr (34). **B.** DIC and fluorescence microscopy images of YPH499 cells transformed with plasmids pEG(KG)-GST-Ø, –IRAK4, –IRAK4(KD), or –IRAK1, following FM 4-64 staining for 75 min. Heterologous protein expression was induced for 8 h in SG Ura^−^ Leu^−^ medium prior to staining. The experiment was carried out with biological triplicates; representative images are shown. Scale bar: 5 μm. The bar graph (right) shows the distribution of vacuolar morphologies across >100 cells per condition, classified according to the scheme panel in A. Bars represent the mean of biological triplicates; error bars indicate standard deviation (SD). Asterisks (***) denote statistically significant differences (p < 0.001, Tukey’s HSD test) for the class B *vps* phenotype.

### *S. cerevisiae* as a sensitive model to assay IRAK4 inhibition: development of a fluorescence-based reporter

Given that human IRAK4 expression causes marked growth inhibition in *S. cerevisiae*, an effect dependent on its kinase activity, we hypothesized that this phenotype could serve as the basis for an in vivo screening platform to identify IRAK4 inhibitors. As a proof of principle, we tested PF-06650833, a well-characterized and highly specific IRAK4 kinase inhibitor (36).

In broth assays, PF-06650833 at concentrations around 1 µM restored the growth of IRAK4-expressing yeast cells to levels comparable to those expressing the catalytically inactive IRAK4(KD) variant (**Fig. 6A**) In solid medium, using a reverse halo assay, spotting 10 mM PF-06650833 onto cellulose discs produced a growth halo exceeding 62 mm in diameter (**Fig. 6B**), indicating diffusion and robust growth rescue. Importantly, no cytotoxicity on yeast cells was observed for PF-06650833 at the tested concentrations. These results support the use of *S. cerevisiae* as a simple, cost-effective, and genetically tractable *in vivo* platform for evaluating specific IRAK4 kinase inhibitors.

**Figure 6.**
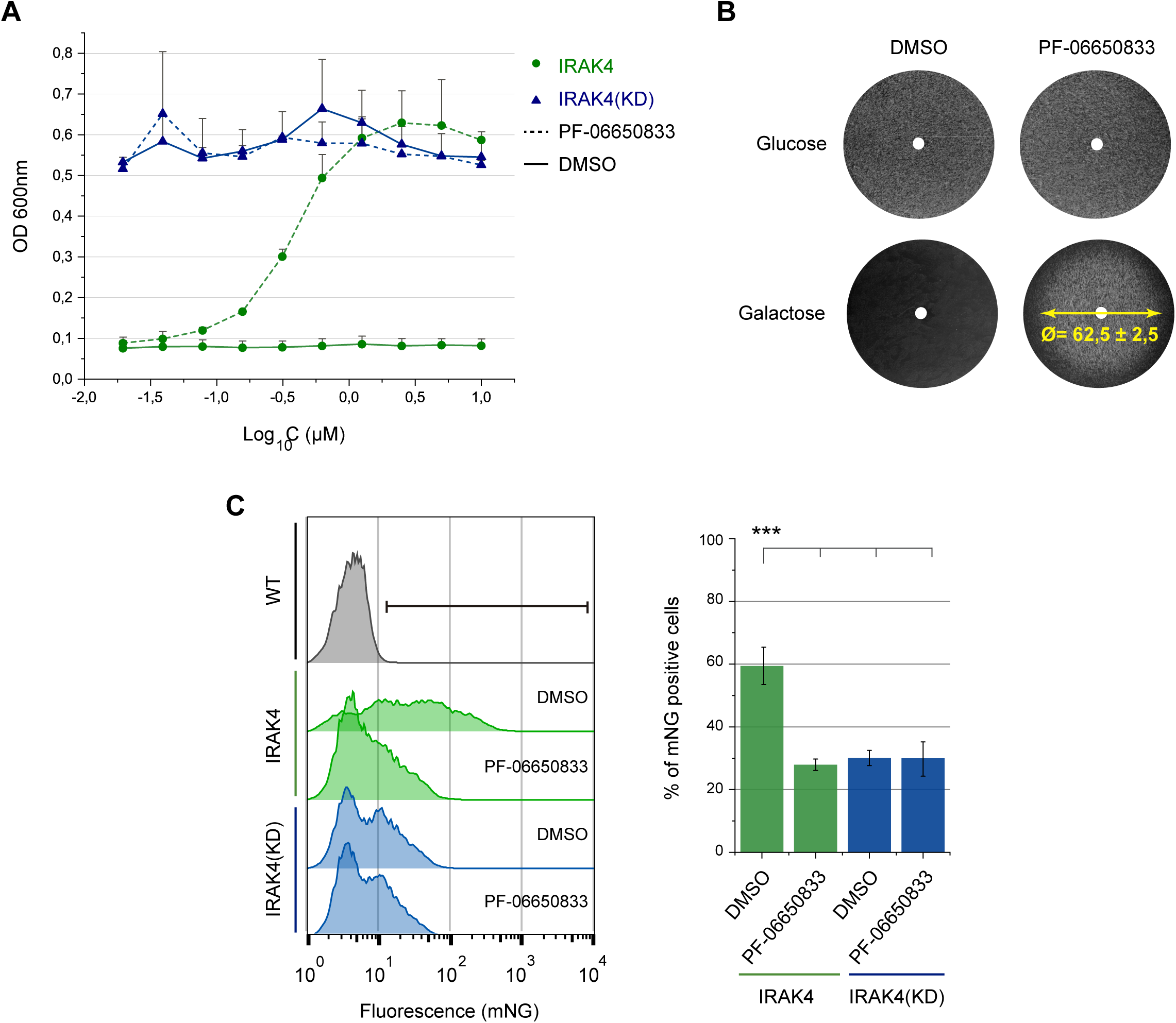
Humanized yeast expressing IRAK4 serves as a platform for screening kinase activity inhibitors. **A.** Growth inhibition assay in liquid medium using PF-06650833, a specific inhibitor of IRAK4 kinase activity. YPH499 cells transformed with pEG(KG)-GST-IRAK4 or pEG(KG)-GST-IRAK4(KD) were cultured in SG Ura^−^ Leu^−^ medium supplemented with serial dilutions of PF-06650833 (from 10 to 0.01953 μM) or equivalent dilutions of the vehicle (DMSO). Absorbance at 600 nm (OD_600_) was measured after 72 h. The graph shows the arithmetic mean of biological triplicates; error bars represent the standard deviation (SD). The x-axis corresponds to the log10 of PF-06650833 concentrations. **B.** Reverse halo assay to evaluate the inhibitory effect of PF-06650833 on IRAK4 catalytic activity in solid medium. YPH499 cells transformed with pEG(KG)-GST-IRAK4 were seeded on SD and SG Ura^−^ Leu^−^ agar plates. Cellulose discs impregnated with 5 mM PF-06650833 or DMSO were placed in the center of the plates. After 72 h of incubation, the diameter of the growth recovery halo was measured. A representative result from three biological triplicates is shown. **C.** Flow cytometry analysis of EVY23 cells (GRE1-mNeonGreen) transformed with pEG(KG)-GST-IRAK4 or pEG(KG)-GST-IRAK4(KD) after 6 h of induction in SG Ura^−^ Leu^−^ medium. PF-06650833 was added at a final concentration of 5 μM at the onset of heterologous protein expression. Left panel: overlaid histogram showing the fluorescence intensity of the Gre1-mNG protein on the x-axis and the number of cellular events on the y-axis. The experiment was performed in biological triplicates for each experimental condition, and a representative result is shown. Right panel: bar graph showing the mean percentage of cells positive for the green fluorescence signal in each condition, based on the threshold established in the histogram for the wild-type strain (WT). Error bars represent the SD. Asterisks (***) indicate a p <0.001, calculated using Tukey’s HSD test.

To enable a scalable, fluorescence-based assay suitable for high-throughput screening, we leveraged our transcriptomic data to develop a transcriptional reporter of IRAK4 activity in yeast. We selected *GRE1*, a hydrophilin-encoding gene that was strongly upregulated in IRAK4-expressing cells but remained unaltered in IRAK4(KD) controls. A reporter construct was generated by fusing the *GRE1* gene to the fluorescent reporter mNeonGreen. When expressed in this background, IRAK4 induced a clear fluorescent signal detectable by flow cytometry, while IRAK4(KD) failed to do so. Moreover, treatment with 5 µM PF-06650833, reduced fluorescence in IRAK4-expressing cells to baseline levels observed in IRAK4(KD) controls (**Fig. 6C**), demonstrating the specificity and responsiveness of the assay to kinase inhibition.

## DISCUSSION

IRAK kinases are essential components of the myddosome, the SMOC that mediates TLR and IL-1R signaling in innate immunity and inflammation (1, 5, 10). Deregulation of these pathways is associated with a range of autoimmune, degenerative and oncogenic disorders (12, 37, 38). In this study, we describe for the first time the heterologous expression of human IRAK kinases in the model organism *S. cerevisiae* and examine their impact on yeast physiology. Together with previous findings on other myddosome components (15, 39), our results lay the groundwork for future synthetic biology approaches aimed at dissecting human signalosomes.

Our transcriptomic, biochemical and phenotypic data suggest that both IRAK1 and IRAK4 function as constitutively active kinases in yeast, showing distinct effects on cell physiology likely driven by differential phosphorylation of endogenous yeast targets. In contrast, IRAK2 showed a minimal impact in our model. Although some kinase activity has been reported for IRAK2, it lacks a canonical catalytic Asp residue and is thus considered an atypical kinase (40, 41). Since each kinase was studied individually, our results support the idea that IRAK4 and IRAK1 undergo autoactivation, while IRAK2 likely requires trans-activation by IRAK4 (42).

Most known phosphorylation events by IRAKs occur within the myddosome environment, and are linked to its assembly, hierarchy, and dynamics (20, 43, 44), with few known substrates outside this signalosome core. The lack of toxicity of kinase-dead IRAK4 [IRAK4(KD)], combined with the strong rescue of IRAK4-induced growth inhibition by the selective inhibitor PF-06650833, confirms that toxicity in yeast is strictly dependent on IRAK4 kinase activity. Therefore, this yeast model could be used to evaluate the functional consequences of IRAK mutations —including alternative splicing variants and disease-associated mutations— on kinase activity. Loss-of-function mutations in *IRAK4* are associated with increased susceptibility to bacterial infections (45, 46), while gain-of-function mutations in both *IRAK1* and *IRAK4* are linked to oncogenesis (47–49). As shown for other human oncoproteins (50, 51), their impact on yeast physiology could serve as a convenient *in vivo* readout for their kinase function.

Growing evidence highlights the interplay between innate immune signaling pathways and immunometabolism (52). TLR/IL-1R signaling via the myddosome has been shown to profoundly influence cellular metabolism (53, 54). Among the most prominent metabolic shifts in activated immune cells is the increase in glycolytic flux (55). In dendritic cells, this response is mediated by TBK1/IKK-ε kinases downstream of TLRs (56), while in LPS-activated macrophages, myddosome assembly was shown critical to enhance glycolysis (57). Similarly, yeast cells reprogram metabolism in response to stress (58, 59). Notably, most transcriptional changes induced by IRAK1 and IRAK4 in yeast were related to metabolic regulation and resembled a nutrient starvation response. Interestingly, although IRAK1 exerted stronger effects on the transcriptome, IRAK4 had a greater inhibitory impact on yeast growth. Both kinases upregulated glycolysis– and glycogen synthesis-related genes, but differed in other metabolic effects: fatty acid β-oxidation was underrepresented in IRAK4-(independently of its kinase activity), yet overrepresented in IRAK1-expressing yeast cells. Additionally, IRAK4 strongly upregulated genes involved in trehalose biosynthesis, typically associated with low glucose perception (60). Since our experiments were performed during exponential growth in galactose (to induce *GAL1*-driven expression) promoter, these changes reflect IRAK-dependent metabolic rewiring rather than carbon source limitation.

Most knowledge on immunometabolism derives from immortalized mouse bone marrow-derived macrophages (iBMDM), which may not fully represent the metabolic features of tissue-resident or human macrophages (61, 62). Thus, alternative models such as yeast could provide insight into the specific properties of human signaling components. Core metabolic pathways and their regulation are highly conserved from yeast to humans. Based on our findings, save the evolutionary distance, IRAK4 may mimic a pro-inflammatory M1-like metabolic profile —marked by mitochondrial fragmentation and reduced β-oxidation— while IRAK1 may resemble an anti-inflammatory M2-like state, with mitochondrial hyperfusion and preserved oxidative function. Therefore, is tempting to speculate that IRAK kinases modulate metabolism via direct phosphorylation of conserved metabolic enzymes or regulators.

Interestingly, in mouse adipocytes, IL-1 stimulation triggers myddosome translocation to the mitochondrial outer membrane through TOM20 recognition, leading to IRAK2-dependent suppression of oxidative phosphorylation (OXPHOS) and β-oxidation (63). This parallels our observation of mitochondrial remodeling in yeast by IRAK kinases and prior reports of MyD88 DD localization to mitochondria-associated membranes in *S. cerevisiae*(15, 16).

Mild stressors such as nutrient limitation can induce mitochondrial hyperfusion — a state linked to enhanced OXPHOS efficiency and protection from autophagy— in both yeast and mammalian cells (64–66). On the contrary, mitochondrial fragmentation, as observed here in IRAK4-expressing yeast cells, is associated with severe stress, such as long-term nutrient deprivation (67), leading to mitochondrial membrane depolarization and subsequent OXPHOS failure (68), eventually favoring mitophagy (69). These features correlate with actin depolarization and loss of polarized growth in IRAK4-expressing yeast, indicating stronger cellular stress than that imposed by IRAK1.

In yeast, TORC1 is typically active during exponential growth phase and suppresses autophagy (70), whereas AMPK (Snf1 in yeast) is activated under energy stress conditions and repressed during rapid growth and nutrient abundance (71). Thus, IRAK4 expression appears to create a metabolic conflict: simultaneously activating glycogen synthesis, TORC1, and downregulating AMPK. The vacuolar hyperfragmentation observed in IRAK4-expressing yeast cells is consistent with TORC1 hyperactivation, as this complex associates with the vacuole and promotes vacuolar fission, resulting in the fragmentation of this organelle. In contrast, under nutrient starvation, TORC1 is downregulated promoting vacuolar fusion and autophagy (72). However, in IRAK4-expressing yeast cells, autophagy occurs despite TORC1 activation, suggesting uncoupling of these pathways.

Targeting the myddosome is a promising therapeutic strategy for inflammatory, autoimmune, and oncologic diseases (73, 74). Hence, IRAK4 has emerged as a druggable target, with several compounds currently in clinical trials (75). Here, we present two simple yeast-based platforms to assess IRAK4 inhibition *in vivo*: one based on growth restoration and another on suppression of a fluorescent transcriptional reporter. Based on our previous experience, both could be adapted for high-throughput screening (76). As proof of concept, we validated our approach using PF-06650833, a selective IRAK4 kinase inhibitor (36). However, kinase inhibition alone may be insufficient to block disease progression, as scaffolding functions of IRAK4 seem to play a major role (10, 77). New strategies such as PROTACs, which promote proteasomal degradation, can simultaneously block both kinase and scaffolding functions (78, 79). Nevertheless, selectively inhibiting kinase activity remains valuable for fine-tuning TLR and IL-1R signaling.

In conclusion, we have established *S. cerevisiae* as a viable platform for functional analyses of human IRAK1 and IRAK4 kinases, and developed specific tools for evaluating cell-permeable IRAK4 kinase-specific inhibitors.

## MATERIALS AND METHODS

### Strains, media, and growth conditions

The *S. cerevisiae* strain YPH499 (*MATa ura3-52 lys2-801_amber ade2-101_ochre trp1-Δ63 his3-Δ200 leu2-Δ1*) was used in all experiments unless otherwise specified. The BY4741 *gas1*Δ strain (used as a PKA signaling control was obtained from the Saccharomyces Genomic Delete Collection (80). Synthetic dextrose (SD) medium contained 2% glucose, 0.17% yeast nitrogen base without amino acids, 0.5% ammonium sulfate, and 0.12% synthetic amino acid drop-out mix, omitting specific components for plasmid selection. In synthetic galactose (SG) and synthetic raffinose (SR) media, glucose was replaced with 2% (w/v) galactose or 1.5% (w/v) raffinose, respectively. Media were prepared with deionized water (Elix Essential 10 system, Merck, RRID: SCR_001287) and solidified with 2.4% (w/v) agar when required. All media were sterilized by autoclaving at 121 °C for 20 min. *GAL1*-driven protein expression was induced by growing cells in SR medium to mid-exponential phase, then refreshing the cultures to an OD₆₀₀ of 0.3 in SG medium lacking appropriate selection amino acids for 6 h, unless otherwise specified.

*Escherichia coli* DH5α was used for routine molecular biology procedures. *E. coli* cultures were grown in LB medium supplemented, as required, with 100 µg/mL ampicillin, 50 µg/mL kanamycin, or 25 µg/mL chloramphenicol. Yeast and bacterial cultures were incubated at 30 °C and 37 °C, respectively, with shaking at 180 rpm.

### Yeast strains construction

The EVY7 (isogenic to YPH499; *snf1::SNF1-6xMYC-TRP1*) and EVY19 (isogenic to YPH499; *atg13::ATG13-6xMYC-TRP1*) strains, which expresses C-terminally Myc-tagged versions of SNF1 and ATG13, respectively, were generated by integration of a *6xMyc-TRP1* cassette. This cassette, encoding six copies of the Myc epitope (EQKLISEEDL), was amplified by PCR from plasmid pRS304 using primers flanked by sequences homologous to the regions upstream and downstream of the stop codon of each gene. Primer sequences are listed in **Suppl. Table S9.**

Strain EVY24 (isogenic to YPH499; *gre1::GRE1-mNeonGreen-ADH1(t)-HygR*), which expresses the Gre1 protein fused at its C-terminus to the green fluorescent protein mNeonGreen (mNG), was generated by genomic integration of a *mNeonGreen-ADH1(t)-HygR* cassette. This cassette was PCR-amplified from plasmid pAP67 and flanked by sequences homologous to the regions upstream and downstream of the *GRE1* stop codon. Integration was verified by PCR on genomic DNA using a forward primer annealing to the intergenic region upstream of GRE1 and a reverse primer within the mNeonGreen coding sequence. High-efficiency transformation of YPH499 with 45 µL of PCR product enabled genomic integration. Primer sequences are listed in **Suppl. Table S9**.

### Plasmid construction

Standard molecular biology techniques were applied following established protocols. Oligonucleotides and plasmids used in this study are listed in **Suppl. Table S9** and **S10**. Human cDNA for IRAK4, IRAK1, and IRAK2 obtained from clones available in DNASU (DNASU Core Facility (RRID: SCR_012185): IRAK1, cloneID 22341; IRAK2, cloneID 22389; IRAK4, cloneID 22346) served as templates for insert amplification. For restriction enzyme-based cloning, PCR products were first subcloned into the pGEM-T® vector (Promega (RRID: SCR_006724), and the resulting inserts were subsequently cloned into the pEG(KG) yeast expression vector (81). The restriction enzymes employed were *Bam*HI and *Xba*I for IRAK4 and IRAK2, and *Xba*I and *Hin*dIII for IRAK1.

Site-directed mutagenesis was carried out using either the two-step method of Wang and Malcolm (82). Briefly, mutagenic primers were designed according to the QuikChange Site-Directed Mutagenesis kit guidelines and amplified using *Pfu*Turbo DNA polymerase (Agilent Technologies, RRID: SCR_013575). Overlapping PCR, used for fusing DNA fragments into a single amplicon, required at least 18 nucleotides of overlap and was performed using the Advantage HD Polymerase (Takara Bio Inc., RRID: SCR_021372). Primer sequences are listed in **Suppl. Table S9**.

### Spot growth assay

Yeast cultures were grown overnight in SD medium. Each culture was then diluted with sterile water in the first column of a 96-well plate to an OD₆₀₀ of 0.5 in a final volume of 200 µL. From this initial dilution, three consecutive tenfold serial dilutions were prepared by transferring 20 µL into 180 µL of sterile water preloaded in the 2nd, 3rd, and 4th columns. These dilutions were subsequently spotted onto SD and SG agar plates using a Multi-blot VP 407AH replicator (V&P Scientific Inc., San Diego, CA, USA). Plates were incubated at 30 °C for 48-72 hs.

### Cell viability assay

Yeast cultures were incubated overnight in SR medium. The following morning, they were diluted to an OD₆₀₀ of 0.3 in SG medium and incubated for 12 hs. Subsequently, cultures were further diluted to an OD₆₀₀ of 0.0001, and 100 µL of these dilutions were plated onto Petri dishes containing YPD medium. After 24 hs of incubation, colony counts were performed. Viability was normalized to 100% based on the control carrying the empty plasmid.

### Multiwell plate inhibition assay

Overnight yeast cultures were grown in SD medium and then diluted to an OD₆₀₀ of 0.05 in SG medium. A 96-well plate was prepared with 100 µL of SG medium per well, except for the first column, where 200 µL of SG medium containing either 10 µM of the test compound or its vehicle was added. A 1:2 serial dilution was performed across columns 1 to 10. Then, 5 µL of yeast culture was added to wells in columns 1–11. Column 11 served as the no-inhibitor growth control (cells only), while column 12 was the negative control (medium with compound or vehicle, no cells). The assay was carried out in biological triplicates. Plates were incubated for 72 h at 30 °C without shaking. After incubation, cell growth was quantified by measuring absorbance at 595 nm using a 96-well plate spectrophotometer (Bio-Rad Laboratories (RRID: SCR_008426) model 680.

### Reverse halo inhibition assay

Yeast liquid cultures incubated overnight in SD medium were diluted to an OD₆₀₀ of 0.5 in both SD and SG media. Then, 100 µL of each culture were spread onto SD and SG agar plates, respectively. To ensure even cell distribution across the surface, sterile glass beads were used. Next, 6 mm cellulose discs impregnated with 15 µL of the test compound (5 mM concentration) were placed in the middle of each plate. The same procedure was followed using the vehicle alone as a control. Plates were incubated at 30 °C for 72 hs. After incubation, the diameter of the growth recovery halo was measured.

### Iodine/iodide staining for glycogen determination

Iodine/iodide staining was used to assess intracellular glycogen accumulation, following the protocol by García *et al*. (83) with minor modifications. Cultures in exponential phase were adjusted to the same OD_600_, and an equivalent number of cells was collected by centrifugation at 2,500 rpm for 3 min. Pellets were resuspended in 1 mL of water, and 330 µL of each sample were transferred and centrifuged again. Cells were then resuspended in 600 µL of an iodine/potassium iodide solution (0.3% I₂, 0.15% KI) and incubated for 3 min at room temperature in the dark. After centrifugation, the pellets were resuspended in 50 µL of water and transferred to a 96-well plate for immediate scanning.

### Transcriptomic analyses

Global gene expression analyses were performed using GeneChip™ Yeast Genome 2.0 Arrays by Affymetrix (RRID: SCR_007817), which cover nearly 100% of *S. cerevisiae* genes. RNA extraction, cDNA synthesis, labeling, hybridization, scanning, and preliminary data analysis were performed at the Genomics Core Facility at the Complutense University of Madrid (RRID: SCR_011166). Yeast cultures were induced in SG medium for 6 hs, and ∼5 × 10⁷ cells were collected. Total RNA was extracted using the NucleoSpin RNA kit (Macherey-Nagel, Düren, Germany) and quantified by Thermo Scientific NanoDrop Lite Spectrophotometer (RRID: SCR_025369). RNA quality was assessed using the Agilent BioAnalyzer 2100 (RRID: SCR_019715). For microarray analysis, 5 µg of total RNA was reverse-transcribed into double-stranded cDNA, which was then in vitro transcribed into biotin-labeled complementary RNA (cRNA) using Affymetrix kits. Labeled cRNA was chemically fragmented and hybridized (15 µg per array) at 45 °C for 16 hs. Arrays were washed, stained with streptavidin–phycoerythrin, and scanned using the GeneChip 3000 7G Microarray Scanner (RRID: SCR_019341).

Raw data (.CEL files) were processed with the GCRMA algorithm by Bioconductor (RRID:SCR_006442), and expression fold changes (FC) were calculated as the ratio of Tukey-weighted means between experimental and control conditions (empty vector). Genes with FC ≥ 2 or ≤ 0.5 and false discovery rate (FDR)< 0.05 were considered differentially expressed. Statistical analyses were performed using Transcriptome Analysis Console (RRID:SCR_016519), and raw data were deposited in GEO under accession number GSE236671. Functional enrichment of differentially expressed genes was performed using PANTHER v18.0 (RRID:SCR_004869), applying Fisher’s exact test with FDR correction (FDR < 0.05). Analyses included GO Biological Process terms and Reactome pathways, with enrichment expressed as fold change relative to the *S. cerevisiae* genome. Gene clustering and visualization were done using MultiExperiment Viewer (MeV v4.9.0; RRID: SCR 001915), employing Euclidean distance and hierarchical clustering for both genes and samples. Transcription factor enrichment was assessed with Yeast Search for Transcriptional Regulators And Consensus Tracking (YEASTRACT+; RRID: SCR_006076), based on DNA-binding evidence. Results were plotted as log2-transformed expression ratios.

### Immunodetection by Western Blotting

Galactose-induced yeast cultures (20 mL) were collected by centrifugation at 2500 rpm and 4 °C during 3 min, and resuspended in 150 µL of lysis buffer (50 mM Tris-HCl pH 7.5; 10% glycerol; 1% Triton X-100; 0.1% NP-40; 0.1% SDS; 150 mM NaCl; 50 mM NaF; 50 mM β-glycerophosphate; 5 mM EDTA; 5 mM Na_2_P_2_O_7_; 1 mM Na_3_VO_4_) supplemented with 3 mM PMSF, 10 mM DTT, and a protease inhibitor cocktail (1 tablet/10 mL, ThermoFisher Scientific, RRID: SCR_008452). Mechanical lysis was performed using glass beads (ø = 0.75–1 mm) and a FastPrep®-24 system (RRID: SCR_018599) (two 30 s cycles at 5.5 m/s). After centrifugation at 13,000 rpm and 4 °C for 10 min, protein concentration in the supernatant was measured at 280 nm using a Beckman DU 640 spectrophotometer (RRID:SCR_008940). Samples were adjusted to the lowest protein concentration and mixed with 2×SDS-PAGE loading buffer (125 mM Tris-HCl pH 6.8; 25% glycerol; 250 mM DTT; 5% SDS; 0.2% bromophenol blue) in a 1:1 ratio and heated at 99 °C for 5 min.

Polyacrylamide gels were prepared with a 10% separating gel (10% acrylamide-bisacrylamide, 0.38 M Tris-HCl pH 8.8, 0.1% SDS, 0.05% ammonium persulfate [APS], 1.3 µL/mL tetramethylethylenediamine [TEMED]) and a 5% stacking gel (5% acrylamide-bisacrylamide, 0.25 M Tris-HCl pH 6.8, 0.1% SDS, 0.1% APS, 2 µL/mL TEMED). 10 µL of protein samples were loaded into the wells, and molecular weight markers (Abcam RRID: SCR_012931) were included. SDS-PAGE was run at 120 V using standard running buffer (25 mM Tris base; 192.6 mM glycine; 0.1% SDS).

After electrophoresis, proteins were transferred onto a 0.45 µm nitrocellulose membrane (Cytiva Life Sciences Amersham Protran, RRID:SCR_013566) using Mini Trans-Blot Cell tanks (Bio-Rad, RRID: SCR_008426) at 110 V for 1 h with transfer buffer (48 mM Tris base; 38.6 mM glycine; 0.037% SDS; 20% ethanol). The membrane was blocked with 5% skim milk in PBS for 1 h at room temperature with gentle agitation. Primary antibody incubation was performed overnight at 4 °C with agitation, using antibodies diluted in 0.1% PBS-T with 1% skim milk. Then, after five 5-min washes with PBS-T, the membrane was incubated for 1 h at room temperature in the dark with fluorophore-conjugated secondary antibodies diluted in the same buffer. Afterwards, washes were repeated. Protein visualization was conducted with the ChemiDoc MP Imaging System (Bio-Rad, RRID: SCR_008426), and band intensity was quantified using Fiji (RRID: SCR_002285). The antibodies used in this study are listed in **Suppl. Table 11.**

### FM 4-64 Staining

Endocytic membranes were visualized using the FM 4–64 dye (Molecular Probes; RRID: SCR_013318), following the protocol by Vida & Emr (34). Yeast cultures (1.5 mL) were centrifuged at 2,500 rpm for 3 min, and the pellet was resuspended in 200 µL of the same medium. FM 4–64 was added to a final concentration of 2.4 µM. Cells were incubated at 30 °C, 950 rpm, in the dark for 1 h and 15 min to allow dye internalization to the vacuolar membrane. After incubation, cells were washed three times with PBS and examined by fluorescence microscopy.

### Microscopy Techniques and Image Processing

For *in vivo* microscopy, galactose-induced cultures were harvested by centrifugation at 3000 rpm for 1 min. Differential interference contrast (DIC) and fluorescence microscopy were performed using either an Eclipse TE2000U inverted fluorescence microscope (Nikon, RRID: SCR_023161) provided with an Orca C4742-95-12ER camera (Hamamatsu Photonics, RRID: SCR_017105) and HCImage software (Hamamatsu, RRID: SCR_015041); or a Leica DMi8 microscope (RRID: SCR_026672). In the latter, images were processed with LAS X software (Leica, RRID: SCR_013673). Further images processing was performed using ImageJ (RRID: SCR_003070) and Adobe Photoshop (RRID: SCR_014199).

### Flow cytometry and ΔΨm and ROS Determination

For flow cytometric analyses on GFP reporter expression, cultures were induced for 5 h with galactose, and 1 mL of the cell culture was harvested for flow cytometric analyses. Rhodamine 123 (Rd123, Sigma-Aldrich) was used to assess mitochondrial membrane potential (ΔΨm), and dihydrorhodamine 123 (DHR123, Sigma-Aldrich) to measure intracellular ROS levels. For this purposes, 1 mL aliquot of yeast culture was incubated with either Rd123 or DHR123 at a final concentration of 5 µg/mL for 30 min at 30 °C, 950 rpm, in the dark. After incubation, cells were diluted 1:10 in PBS. Flow cytometry experiments were performed using a BD FACScan Scan (RRID: SCR_019596) equipped with a 488 nm excitation laser and emission filters with a bandwidth of 530/30 (FL1), 585/42 (FL2), and 670 (FL3) nm. In all cases, at least 10,000 cells per sample were analyzed, and co-staining with propidium iodide (PI) was performed to exclude fluorescence signal from dead cells. Data were processed using FlowJo (RRID: SCR_008520) software. Analyses were performed at the Flow Cytometry and Fluorescence Microscopy Unit of University of Madrid (RRID: SCR_011166).

### Statistical Analyses

Data were analyzed using Microsoft Excel (RRID: SCR_016137) and IBM SPSS Statistics (RRID: SCR_016479), and graphical representations were generated with Origin (RRID:SCR_014212). Normality was first assessed using the Shapiro-Wilk test. When the null hypothesis (H₀) could not be rejected, a two-tailed unpaired Student’s t-test was used for comparisons between two groups, and One-Way ANOVA followed by Tukey’s Honestly Significant Difference (HSD) *post hoc* test was applied for comparisons among more than two groups. If the null hypothesis (H₀) was rejected and more than two groups were compared, the Kruskal–Wallis test followed by Dunn’s post hoc test with Bonferroni correction was performed. Asterisks in the figures denote statistically significant differences, with *, **, and *** indicating p < 0.05, p < 0.01, and p < 0.001, respectively. When data followed a normal distribution, the arithmetic mean and standard deviation (SD) were calculated; otherwise, the median and interquartile range (IQR, Q3-Q1) were reported.

## ACKNOWLEDGMENTS

This research is part of the I+D+i Grant PID2022-138591NB-I00, funded by MCIN/AEI/10.13039/501100011033 (Ministerio de Ciencia e Innovación, Spain). E.d.V. was supported by a predoctoral contract from Universidad Complutense de Madrid (UCM)-Santander. We are grateful to the staff at the Genomics Core Unit at the Complutense University of Madrid (UCM) for sequencing and help with transcriptomic experiments, and to the UCM Flow Cytometry and Fluorescence Microscopy Unit. We also wish to acknowledge all members of Research Unit 3 at the Dpt. of Microbiology and Parasitology (UCM) for discussion and continuous support.

The authors have no relevant financial or non-financial interests to disclose.

## AUTHOR CONTRIBUTIONS

E.d.V., T. F-A., M.M and V.C. contributed to the design of the study. All material preparation, data collection and analysis were performed by E.d.V., except for the initial IRAK gene cloning, which were contributed by G.G. The first draft of the manuscript was written by E.d.V. and V.C. All authors have read and approved the manuscript.

