## Supplemental Tables, Figures and Methods for "Human IRAK kinases differentially alter metabolic regulation and mitochondrial function in *Saccharomyces cerevisiae*"

Del Val et al.,

### SUPPLEMENTARY TABLES AND FIGURES

**Table S1. Yeast genes differentially expressed after IRAK1 expression**

| Gene name | Fold change | FDR |
|---|---|---|
| **UPREGULATED** | | |
| *DDI2; DDI3 / YFL061W* | **21.78** | 1.84E-09 |
| *FDH1 / YOR388C* | **20.75** | 2.33E-08 |
| *SSA4 / YER103W* | **18.31** | 1.07E-12 |
| *CHA1 / YCL064C* | **18.11** | 8.87E-07 |
| *GRE1 / YPL223C* | **16.66** | 0.0007 |
| *TSA2 / YDR453C* | **15.42** | 0.0004 |
| *YKL107W / YKL107W* | **15.41** | 1.43E-05 |
| *TKL2 / YBR117C* | **15.36** | 1.94E-09 |
| *HEF3 / YNL014W* | **15.07** | 3.76E-06 |
| *YJL045W / YJL045W* | **13.83** | 8.18E-05 |
| *SSA3 / YBL075C* | **13.75** | 1.86E-07 |
| *DMC1 / YER179W* | **13.38** | 1.73E-09 |
| *SPG1 / YGR236C* | **13.02** | 5.45E-10 |
| *HSP32; HSP33; SNO4 / YMR322C* | **11.62** | 7.50E-12 |
| *GIP2 / YER054C* | **11.04** | 5.45E-10 |
| *PMA2 / YPL036W* | **10.64** | 1.86E-07 |
| *PAI3 / YMR174C* | **10.42** | 9.30E-06 |
| *YLR053C / YLR053C* | **9.82** | 3.89E-08 |
| *YLR460C / YLR460C* | **9.69** | 0.0039 |
| *HBT1 / YDL223C* | **9.66** | 7.94E-09 |
| *SHH3 / YMR118C* | **9.57** | 0.0072 |
| *TIR3 / YIL011W* | **9.5** | 0.0001 |
| *YCR102C / YCR102C* | **9.34** | 0.0004 |
| *SIP18 / YMR175W* | **8.89** | 0.0004 |
| *RTN2 / YDL204W* | **8.77** | 2.38E-08 |
| *YJL144W / YJL144W* | **8.31** | 7.08E-09 |
| *YMR018W / YMR018W* | **8.24** | 7.06E-07 |
| *HSP26 / YBR072W* | **8.03** | 1.44E-07 |
| *RCK1 / YGL158W* | **8.03** | 0.0013 |
| *SHH4 / YLR164W* | **7.98** | 9.71E-07 |
| *ALD3 / YMR169C* | **7.84** | 2.24E-06 |
| *BTN2 / YGR142W* | **7.81** | 6.43E-09 |
| *NQM1 / YGR043C* | **7.66** | 1.73E-09 |
| *OYE3 / YPL171C* | **7.55** | 0.0021 |
| *BAG7 / YOR134W* | **7.12** | 2.80E-10 |
| *YML083C / YML083C* | **7.05** | 4.49E-05 |
| *YDR034W-B / YDR034W-B* | **7.05** | 1.88E-05 |
| *CRF1 / YDR223W* | **7.04** | 1.26E-08 |
| *HSP31 / YDR533C* | **7.02** | 0.0006 |

| | | |
|---|---|---|
| *SPS100 / YHR139C* | **6.86** | 0.0001 |
| *PRR2 / YDL214C* | **6.78** | 3.97E-07 |
| *FMP16 / YDR070C* | **6.6** | 3.28E-08 |
| *ERR1; ERR2; ERR3 / YMR323W* | **6.56** | 6.16E-06 |
| *HXT11 / YOL156W* | **6.5** | 7.55E-08 |
| *RGI2 / YIL057C* | **6.48** | 7.08E-09 |
| *RTC3 / YHR087W* | **6.45** | 3.27E-07 |
| *FMP45 / YDL222C* | **6.36** | 3.85E-09 |
| *YGR066C / YGR066C* | **6.31** | 0.0010 |
| *FRE4 / YNR060W* | **6.13** | 1.17E-10 |
| *NDE2 / YDL085W* | **6.04** | 1.82E-09 |
| *UIP4 / YPL186C* | **5.94** | 1.27E-09 |
| *PEX18 / YHR160C* | **5.87** | 5.45E-10 |
| *YPL277C / YPL277C* | **5.84** | 6.51E-08 |
| *PHM7 / YOL084W* | **5.79** | 4.39E-07 |
| *DSF1; YNR073C / YEL070W* | **5.67** | 1.19E-05 |
| *ADY2 / YCR010C* | **5.59** | 5.48E-08 |
| *DCI1 / YOR180C* | **5.58** | 2.40E-07 |
| *SLZ1 / YNL196C* | **5.52** | 1.74E-07 |
| *IMA5 / YJL216C* | **5.38** | 4.24E-08 |
| *LEE1 / YPL054W* | **5.37** | 1.84E-09 |
| *SPO1 / YNL012W* | **5.24** | 3.22E-11 |
| *FMP48 / YGR052W* | **5.24** | 6.73E-09 |
| *CTT1 / YGR088W* | **5.18** | 3.22E-08 |
| *YNL195C / YNL195C* | **5.06** | 1.77E-09 |
| *YMR090W / YMR090W* | **5.03** | 4.16E-05 |
| *SOL4 / YGR248W* | **4.86** | 1.84E-09 |
| *HXT9 / YJL219W* | **4.85** | 4.84E-06 |
| *GND2 / YGR256W* | **4.8** | 4.39E-05 |
| *YDL218W / YDL218W* | **4.73** | 1.76E-05 |
| *MSC1 / YML128C* | **4.73** | 5.45E-10 |
| *HPF1 / YIL169C* | **4.71** | 2.53E-05 |
| *IMA1 / YGR287C* | **4.67** | 9.77E-06 |
| *IME1 / YJR094C* | **4.66** | 1.81E-08 |
| *PIR3 / YKL163W* | **4.64** | 2.12E-06 |
| *YPL278C / YPL278C* | **4.57** | 5.70E-08 |
| *IDP3 / YNL009W* | **4.55** | 4.04E-08 |
| *HSP42 / YDR171W* | **4.55** | 7.92E-10 |
| *SPI1 / YER150W* | **4.53** | 1.50E-08 |
| *YLR346C / YLR346C* | **4.48** | 0.0038 |
| *REG2 / YBR050C* | **4.48** | 5.45E-10 |
| *SUL1 / YBR294W* | **4.47** | 2.80E-06 |
| *PHO92 / YDR374C* | **4.38** | 0.0023 |
| *BIO5 / YNR056C* | **4.28** | 2.40E-07 |
| *POT1 / YIL160C* | **4.25** | 1.09E-07 |

| | | |
|---|---|---|
| *YMR034C / YMR034C* | **4.22** | 4.11E-09 |
| *GRE2 / YOL151W* | **4.21** | 6.84E-06 |
| *MIG3 / YER028C* | **4.21** | 0.0002 |
| *MPO1 / YGL010W* | **4.13** | 3.64E-10 |
| *IMA3 / YIL172C* | **4.12** | 4.05E-09 |
| *HXT8 / YJL214W* | **4.06** | 7.20E-07 |
| *CUR1 / YPR158W* | **4.05** | 8.41E-09 |
| *YJR096W / YJR096W* | **4.03** | 5.04E-08 |
| *GPX1 / YKL026C* | **3.97** | 2.66E-09 |
| *HPF1 / YOL155C* | **3.96** | 0.0002 |
| *FUS1 / YCL027W* | **3.96** | 4.05E-05 |
| *YEL057C / YEL057C* | **3.95** | 1.62E-08 |
| *OPI10 / YOL032W* | **3.94** | 6.43E-09 |
| *YML131W / YML131W* | **3.92** | 7.48E-07 |
| *YHR138C / YHR138C* | **3.88** | 1.22E-07 |
| *PXA2 / YKL188C* | **3.85** | 7.08E-09 |
| *VPS73 / YGL104C* | **3.84** | 2.67E-09 |
| *GAD1 / YMR250W* | **3.84** | 2.86E-07 |
| *YNL134C / YNL134C* | **3.84** | 0.0005 |
| *FRT2 / YAL028W* | **3.84** | 5.45E-10 |
| *DCS2 / YOR173W* | **3.83** | 1.64E-09 |
| *DBP1 / YPL119C* | **3.82** | 4.71E-08 |
| *TDH1 / YJL052W* | **3.82** | 0.0002 |
| *YER053C-A / YER053C-A* | **3.79** | 1.22E-07 |
| *ECM4 / YKR076W* | **3.78** | 5.16E-09 |
| *SPO20 / YMR017W* | **3.77** | 9.58E-09 |
| *YMR147W / YMR147W* | **3.76** | 1.06E-06 |
| *SPG4 / YMR107W* | **3.75** | 6.86E-08 |
| *YOR186W / YOR186W* | **3.73** | 0.0025 |
| *GPG1 / YGL121C* | **3.73** | 4.39E-07 |
| *YOL131W / YOL131W* | **3.73** | 0.0014 |
| *SPL2 / YHR136C* | **3.68** | 2.46E-05 |
| *YGR201C / YGR201C* | **3.65** | 0.0004 |
| *YKL050C / YKL050C* | **3.62** | 0.0023 |
| *ECI1 / YLR284C* | **3.59** | 1.41E-07 |
| *YLR031W / YLR031W* | **3.57** | 3.38E-05 |
| *ICS2 / YBR157C* | **3.56** | 1.88E-08 |
| *YNL194C / YNL194C* | **3.51** | 1.76E-06 |
| *RTS3 / YGR161C* | **3.51** | 1.41E-07 |
| *SAF1 / YBR280C* | **3.5** | 7.42E-09 |
| *ECM8 / YBR076W* | **3.49** | 5.23E-06 |
| *SRL4 / YPL033C* | **3.47** | 0.0322 |
| *RRT12 / YCR045C* | **3.42** | 0.0108 |
| *CTA1 / YDR256C* | **3.42** | 1.71E-07 |
| *YNR034W-A / YNR034W-A* | **3.41** | 0.0002 |

| | | |
|---|---|---|
| *GSY1 / YFR015C* | **3.39** | 9.58E-10 |
| *STF2 / YGR008C* | **3.38** | 1.77E-09 |
| *SHC1 / YER096W* | **3.37** | 0.0014 |
| *XBP1 / YIL101C* | **3.34** | 2.10E-07 |
| *HOP1 / YIL072W* | **3.33** | 1.22E-07 |
| *PAU21 / YOR394W* | **3.3** | 1.35E-07 |
| *YNL200C / YNL200C* | **3.29** | 4.17E-07 |
| *YGR127W / YGR127W* | **3.28** | 6.92E-08 |
| *CRG1 / YHR209W* | **3.28** | 2.56E-08 |
| *BDH2 / YAL061W* | **3.27** | 3.51E-08 |
| *CRS5 / YOR031W* | **3.27** | 0.0004 |
| *DIA3 / YDL024C* | **3.27** | 9.77E-06 |
| *SSE2 / YBR169C* | **3.27** | 6.01E-08 |
| *YPL119C-A / YPL119C-A* | **3.22** | 6.14E-07 |
| *GLG1 / YKR058W* | **3.2** | 9.79E-08 |
| *ATG34 / YOL083W* | **3.17** | 1.50E-08 |
| *MOH1 / YBL049W* | **3.17** | 7.30E-06 |
| *ALP1 / YNL270C* | **3.16** | 2.06E-05 |
| *YLR415C / YLR415C* | **3.14** | 1.44E-05 |
| *MIH1 / YMR036C* | **3.13** | 6.27E-09 |
| *YMR196W / YMR196W* | **3.13** | 2.72E-07 |
| *YBR285W / YBR285W* | **3.13** | 2.12E-07 |
| *ATG8 / YBL078C* | **3.12** | 8.17E-07 |
| *CIT3 / YPR001W* | **3.08** | 5.74E-06 |
| *TFS1 / YLR178C* | **3.08** | 1.92E-08 |
| *YOR389W / YOR389W* | **3.06** | 3.01E-06 |
| *GPH1 / YPR160W* | **3.05** | 1.19E-08 |
| *PXA1 / YPL147W* | **3.02** | 3.89E-08 |
| *YJL133C-A / YJL133C-A* | **3.02** | 8.71E-09 |
| *ATG29 / YPL166W* | **2.98** | 0.0001 |
| *SET4 / YJL105W* | **2.98** | 8.45E-05 |
| *POX1 / YGL205W* | **2.97** | 2.07E-06 |
| *ETR1 / YBR026C* | **2.97** | 1.04E-08 |
| *YJR115W / YJR115W* | **2.97** | 9.70E-07 |
| *YGR174W-A / YGR174W-A* | **2.96** | 0.0001 |
| *EMP46 / YLR080W* | **2.96** | 2.83E-07 |
| *PHM8 / YER037W* | **2.91** | 0.0065 |
| *GLC3 / YEL011W* | **2.9** | 3.51E-08 |
| *STR3 / YGL184C* | **2.9** | 0.0044 |
| *ALD2 / YMR170C* | **2.9** | 1.58E-07 |
| *PUT1 / YLR142W* | **2.88** | 0.0003 |
| *YJL163C / YJL163C* | **2.88** | 1.32E-07 |
| *YKL151C / YKL151C* | **2.85** | 6.14E-05 |
| *YPL113C / YPL113C* | **2.85** | 2.99E-06 |
| *ATG9 / YDL149W* | **2.84** | 8.21E-08 |

| | | |
|---|---|---|
| *YLR030W / YLR030W* | **2.83** | 1.27E-05 |
| *OM45 / YIL136W* | **2.78** | 9.34E-07 |
| *MBF1 / YOR298C-A* | **2.77** | 3.02E-08 |
| *PDR10 / YOR328W* | **2.76** | 0.0001 |
| *YET2 / YMR040W* | **2.75** | 0.0002 |
| *YPR015C / YPR015C* | **2.75** | 0.0002 |
| *XYL2 / YLR070C* | **2.73** | 8.80E-08 |
| *TMA17 / YDL110C* | **2.72** | 2.39E-08 |
| *MHO1 / YJR008W* | **2.72** | 2.87E-09 |
| *YBR184W / YBR184W* | **2.71** | 2.40E-05 |
| *AMS1 / YGL156W* | **2.71** | 1.97E-08 |
| *YNL144C / YNL144C* | **2.71** | 1.86E-07 |
| *CTF3 / YLR381W* | **2.71** | 1.41E-06 |
| *TMA10 / YLR327C* | **2.7** | 5.76E-07 |
| *ATH1 / YPR026W* | **2.7** | 6.51E-08 |
| *PGM2 / YMR105C* | **2.7** | 6.61E-09 |
| *HSP82 / YPL240C* | **2.69** | 1.66E-07 |
| *GLO4 / YOR040W* | **2.69** | 0.0002 |
| *YLR446W / YLR446W* | **2.69** | 6.36E-07 |
| *ARA2 / YMR041C* | **2.68** | 7.08E-09 |
| *YGL081W / YGL081W* | **2.68** | 6.95E-06 |
| *HXT5 / YHR096C* | **2.68** | 4.49E-06 |
| *WSC4 / YHL028W* | **2.67** | 7.32E-07 |
| *RNY1 / YPL123C* | **2.65** | 4.59E-08 |
| *GDH3 / YAL062W* | **2.65** | 0.0004 |
| *FOX2 / YKR009C* | **2.65** | 1.35E-06 |
| *YEL020C / YEL020C* | **2.64** | 3.15E-05 |
| *YHL044W / YHL044W* | **2.64** | 1.24E-06 |
| *FES1 / YBR101C* | **2.64** | 3.33E-08 |
| *GSY2 / YLR258W* | **2.64** | 3.20E-07 |
| *37226 / YLR348C* | **2.64** | 2.39E-05 |
| *REC104 / YHR157W* | **2.62** | 3.26E-05 |
| *SYM1 / YLR251W* | **2.62** | 6.48E-08 |
| *YHR097C / YHR097C* | **2.62** | 2.20E-08 |
| *SOL1 / YNR034W* | **2.61** | 3.09E-06 |
| *PCD1 / YLR151C* | **2.6** | 1.56E-05 |
| *YLL056C / YLL056C* | **2.6** | 6.89E-05 |
| *YMR206W / YMR206W* | **2.6** | 3.38E-07 |
| *RAD51 / YER095W* | **2.6** | 1.69E-07 |
| *YKL162C / YKL162C* | **2.6** | 1.69E-07 |
| *SDS24 / YBR214W* | **2.6** | 2.68E-07 |
| *PRB1 / YEL060C* | **2.59** | 6.73E-09 |
| *YGR053C / YGR053C* | **2.58** | 1.46E-05 |
| *COX5B / YIL111W* | **2.57** | 7.12E-07 |
| *CTL1 / YMR180C* | **2.56** | 2.66E-09 |

| | | |
|---|---|---|
| *RIM4 / YHL024W* | **2.55** | 9.01E-07 |
| *YNR064C / YNR064C* | **2.55** | 1.16E-06 |
| *SIS1 / YNL007C* | **2.55** | 7.44E-07 |
| *GAP1 / YKR039W* | **2.55** | 1.00E-06 |
| *APJ1 / YNL077W* | **2.55** | 7.15E-08 |
| *YGR174W-A / YGR174W-A* | **2.54** | 0.0006 |
| *KIN82 / YCR091W* | **2.53** | 3.20E-06 |
| *YLR149C / YLR149C* | **2.53** | 7.14E-07 |
| *MXR1 / YER042W* | **2.53** | 3.56E-06 |
| *YSP3 / YOR003W* | **2.53** | 8.53E-06 |
| *SGA1 / YIL099W* | **2.52** | 0.0140 |
| *TDA10 / YGR205W* | **2.52** | 1.80E-05 |
| *TRR2 / YHR106W* | **2.52** | 6.92E-08 |
| *PAU3 / YCR104W* | **2.51** | 4.04E-08 |
| *UGX2 / YDL169C* | **2.51** | 2.25E-07 |
| *YPT53 / YNL093W* | **2.49** | 2.36E-06 |
| *YPL014W / YPL014W* | **2.48** | 0.0003 |
| *YPT35 / YHR105W* | **2.47** | 8.26E-05 |
| *HUL4 / YJR036C* | **2.47** | 3.80E-07 |
| *FAT3 / YKL187C* | **2.46** | 6.50E-07 |
| *ENO1 / YGR254W* | **2.46** | 4.05E-05 |
| *YMR175W-A / YMR175W-A* | **2.45** | 0.0218 |
| *SSU1 / YPL092W* | **2.45** | 1.69E-07 |
| *UBP13 / YBL067C* | **2.44** | 9.43E-08 |
| *URA10 / YMR271C* | **2.44** | 1.16E-05 |
| *YEF1 / YEL041W* | **2.44** | 9.06E-06 |
| *ACS1 / YAL054C* | **2.43** | 9.71E-07 |
| *YFL054C / YFL054C* | **2.43** | 1.22E-07 |
| *YDL199C / YDL199C* | **2.42** | 3.42E-06 |
| *IKS1 / YJL057C* | **2.42** | 4.58E-07 |
| *YMR114C / YMR114C* | **2.41** | 6.41E-07 |
| *YGP1 / YNL160W* | **2.41** | 4.59E-08 |
| *DSD1 / YGL196W* | **2.4** | 4.13E-07 |
| *HSP104 / YLL026W* | **2.4** | 7.90E-08 |
| *IGD1 / YFR017C* | **2.39** | 6.51E-08 |
| *YBR241C / YBR241C* | **2.38** | 9.43E-08 |
| *FUN19 / YAL034C* | **2.38** | 4.24E-07 |
| *YPL277C / YPL277C* | **2.38** | 0.0004 |
| *FAA2 / YER015W* | **2.38** | 1.06E-05 |
| *PIR5 / YJL160C* | **2.37** | 1.41E-06 |
| *ZTA1 / YBR046C* | **2.37** | 9.38E-08 |
| *GSP2 / YOR185C* | **2.36** | 3.28E-05 |
| *YNL115C / YNL115C* | **2.36** | 1.32E-07 |
| *YMR158C-A / YMR158C-A* | **2.36** | 0.0002 |
| *PIG1 / YLR273C* | **2.36** | 9.45E-08 |

| | | |
|---|---|---|
| *PFK26 / YIL107C* | **2.35** | 6.56E-08 |
| *IGO2 / YHR132W-A* | **2.35** | 1.56E-07 |
| *HXK1 / YFR053C* | **2.35** | 2.10E-07 |
| *YDR003W-A / YDR003W-A* | **2.35** | 1.32E-05 |
| *ECM11 / YDR446W* | **2.34** | 0.0003 |
| *SPS19 / YNL202W* | **2.33** | 1.85E-06 |
| *YER039C-A / YER039C-A* | **2.33** | 1.83E-05 |
| *MET28 / YIR017C* | **2.33** | 1.51E-05 |
| *ARG82 / YDR173C* | **2.33** | 5.63E-05 |
| *FYV10 / YIL097W* | **2.33** | 1.26E-06 |
| *SPG5 / YMR191W* | **2.32** | 1.81E-08 |
| *YOR289W / YOR289W* | **2.31** | 9.50E-06 |
| *YIL055C / YIL055C* | **2.31** | 3.78E-08 |
| *DDR2 / YOL052C-A* | **2.31** | 1.68E-06 |
| *ECL1 / YGR146C* | **2.29** | 4.13E-07 |
| *OXR1 / YPL196W* | **2.29** | 1.09E-07 |
| *YIR014W / YIR014W* | **2.29** | 1.69E-07 |
| *GIP1 / YBR045C* | **2.29** | 1.79E-06 |
| *AIM19 / YIL087C* | **2.28** | 2.88E-06 |
| *YLR108C / YLR108C* | **2.28** | 0.0001 |
| *SDP1 / YIL113W* | **2.28** | 4.70E-05 |
| *YCT1 / YLL055W* | **2.28** | 0.0053 |
| *REE1 / YJL217W* | **2.27** | 2.35E-07 |
| *AAC1 / YMR056C* | **2.27** | 1.85E-05 |
| *PRM8 / YGL053W* | **2.27** | 0.0012 |
| *BRE2 / YLR015W* | **2.27** | 5.76E-06 |
| *YHR140W / YHR140W* | **2.27** | 3.12E-07 |
| *YIL169C / YIL169C* | **2.27** | 2.90E-05 |
| *LSO1 / YJR005C-A* | **2.27** | 0.0026 |
| *YDL124W / YDL124W* | **2.26** | 3.61E-08 |
| *OSW5 / YMR148W* | **2.25** | 1.90E-06 |
| *YBL039W-B / YBL039W-A* | **2.25** | 5.98E-06 |
| *RNR3 / YIL066C* | **2.25** | 6.57E-06 |
| *GAC1 / YOR178C* | **2.25** | 3.98E-07 |
| *GTO1 / YGR154C* | **2.25** | 0.0004 |
| *ATG31 / YDR022C* | **2.25** | 1.36E-06 |
| *PDR18 / YNR070W* | **2.24** | 3.62E-06 |
| *VHS1 / YDR247W* | **2.24** | 5.98E-06 |
| *TPK1 / YJL164C* | **2.24** | 6.85E-08 |
| *GTT1 / YIR038C* | **2.23** | 2.50E-05 |
| *YDR018C / YDR018C* | **2.23** | 7.14E-06 |
| *TPS1 / YBR126C* | **2.23** | 7.19E-06 |
| *NRG1 / YDR043C* | **2.23** | 1.36E-07 |
| *RGI1 / YER067W* | **2.22** | 1.84E-05 |
| *CPR6 / YLR216C* | **2.22** | 2.23E-08 |

| | | |
|---|---|---|
| *RAD34 / YDR314C* | **2.22** | 0.0001 |
| *RTK1 / YDL025C* | **2.22** | 2.10E-07 |
| *MPC1 / YGL080W* | **2.22** | 9.20E-06 |
| *SRX1 / YKL086W* | **2.21** | 0.0023 |
| *STI1 / YOR027W* | **2.2** | 1.71E-07 |
| *ALT2 / YDR111C* | **2.2** | 3.62E-06 |
| *KSP1 / YHR082C* | **2.19** | 2.01E-05 |
| *YGL036W / YGL036W* | **2.19** | 2.41E-05 |
| *GDB1 / YPR184W* | **2.18** | 6.22E-06 |
| *PEX11 / YOL147C* | **2.18** | 2.11E-06 |
| *MEI4 / YER044C-A* | **2.18** | 0.0002 |
| *VHS3 / YOR054C* | **2.17** | 3.66E-07 |
| *FMP52 / YER004W* | **2.17** | 1.82E-06 |
| *HEM14 / YER014W* | **2.17** | 1.96E-07 |
| *HFD1 / YMR110C* | **2.16** | 1.93E-07 |
| *NIP100 / YPL174C* | **2.16** | 2.64E-05 |
| *PIN3 / YPR154W* | **2.15** | 1.86E-05 |
| *YLR297W / YLR297W* | **2.15** | 2.74E-06 |
| *CYC7 / YEL039C* | **2.15** | 5.19E-05 |
| *YLR173W / YLR173W* | **2.15** | 3.33E-08 |
| *ATG16 / YMR159C* | **2.14** | 0.0028 |
| *AMA1 / YGR225W* | **2.14** | 0.0002 |
| *NGR1 / YBR212W* | **2.14** | 8.56E-05 |
| *YBR056W / YBR056W* | **2.14** | 4.82E-07 |
| *TSL1 / YML100W* | **2.14** | 3.32E-06 |
| *THI20 / YOL055C* | **2.14** | 0.0001 |
| *MND2 / YIR025W* | **2.14** | 0.0001 |
| *GYP7 / YDL234C* | **2.13** | 1.17E-06 |
| *STB2 / YMR053C* | **2.13** | 6.01E-08 |
| *MGA1 / YGR249W* | **2.12** | 3.68E-05 |
| *FUN14 / YAL008W* | **2.11** | 2.22E-05 |
| *PDR11 / YIL013C* | **2.11** | 1.17E-06 |
| *YKL071W / YKL071W* | **2.11** | 0.0008 |
| *TR / YLL060C* | **2.11** | 0.0023 |
| *YPR127W / YPR127W* | **2.1** | 0.0032 |
| *ATG14 / YBR128C* | **2.1** | 1.69E-07 |
| *HPA2 / YPR193C* | **2.1** | 0.0002 |
| *CSM4 / YPL200W* | **2.1** | 0.0005 |
| *PCS60 / YBR222C* | **2.1** | 0.0037 |
| *YKL133C / YKL133C* | **2.09** | 3.43E-06 |
| *ATG1 / YGL180W* | **2.09** | 5.04E-07 |
| *PHO5 / YBR093C* | **2.09** | 2.28E-05 |
| *RPN4 / YDL020C* | **2.09** | 5.42E-07 |
| *MSS11 / YMR164C* | **2.08** | 3.68E-05 |
| *ATG4 / YNL223W* | **2.08** | 1.01E-05 |

| | | |
|---|---|---|
| *SFA1 / YDL168W* | **2.08** | 6.17E-06 |
| *RTS2 / YOR077W* | **2.07** | 6.85E-05 |
| *PCH2 / YBR186W* | **2.07** | 0.0001 |
| *ATG7 / YHR171W* | **2.07** | 2.29E-06 |
| *SGT2 / YOR007C* | **2.06** | 1.35E-06 |
| *IRC15 / YPL017C* | **2.06** | 1.72E-05 |
| *YLR125W / YLR125W* | **2.06** | 1.34E-05 |
| *YMR181C / YMR181C* | **2.06** | 9.62E-06 |
| *PPE1 / YHR075C* | **2.06** | 6.79E-07 |
| *BNS1 / YGR230W* | **2.06** | 5.67E-07 |
| *LAM5 / YFL042C* | **2.05** | 2.24E-06 |
| *TOS8 / YGL096W* | **2.05** | 0.0003 |
| *YHL042W / YHL042W* | **2.05** | 3.81E-05 |
| *UGA1 / YGR019W* | **2.05** | 3.62E-06 |
| *VTI1 / YMR197C* | **2.05** | 3.94E-05 |
| *TLG2 / YOL018C* | **2.04** | 6.44E-05 |
| *PTC6 / YCR079W* | **2.04** | 1.78E-07 |
| *VID30 / YGL227W* | **2.04** | 5.67E-07 |
| *MPS2 / YGL075C* | **2.04** | 0.0020 |
| *POM33 / YLL023C* | **2.03** | 2.40E-07 |
| *TEN1 / YLR010C* | **2.03** | 7.23E-06 |
| *COQ4 / YDR204W* | **2.03** | 3.97E-07 |
| *OPT1 / YJL212C* | **2.03** | 0.0016 |
| *MPC54 / YOR177C* | **2.02** | 0.0011 |
| *MLH3 / YPL164C* | **2.02** | 8.39E-07 |
| *PLB2 / YMR006C* | **2.02** | 1.12E-05 |
| *YMR084W / YMR084W* | **2.01** | 0.0001 |
| *YRR1 / YOR162C* | **2.01** | 4.13E-07 |
| *SUE1 / YPR151C* | **2.01** | 2.54E-06 |
| *MPC3 / YGR243W* | **2.01** | 1.64E-06 |
| *YMR085W / YMR085W* | **2.01** | 1.33E-06 |
| *BSC5 / YNR069C* | **2** | 0.0002 |
| *YMR160W / YMR160W* | **2** | 1.42E-07 |
| *PIG2 / YIL045W* | **2** | 3.63E-05 |

| DOWNREGULATED | | |
|---|---|---|
| *YGR079W / YGR079W* | **0.50** | 0.0002 |
| *SRB2 / YHR041C* | **0.50** | 0.0006 |
| *AKR1 / YDR264C* | **0.50** | 0.0021 |
| *GIN4 / YDR507C* | **0.50** | 2.10E-07 |
| *EFG1 / YGR272C* | **0.50** | 0.0032 |
| *YGL230C / YGL230C* | **0.50** | 5.01E-06 |
| *CDC60 / YPL160W* | **0.50** | 3.61E-08 |
| *RPL6B / YLR448W* | **0.50** | 3.51E-06 |
| *RPL2A; RPL2B / YIL018W* | **0.49** | 3.80E-05 |
| *PRM7 / YDL039C* | **0.49** | 0.0028 |
| *MRS3 / YJL133W* | **0.49** | 4.66E-07 |
| *IMD2 / YHR216W* | **0.49** | 0.0019 |
| *ATP5 / YDR298C* | **0.49** | 0.0035 |
| *GDH1 / YOR375C* | **0.49** | 4.24E-08 |
| *MLS1 / YNL117W* | **0.49** | 8.43E-07 |
| *HEM3 / YDL205C* | **0.49** | 0.0001 |
| *FRE1 / YLR214W* | **0.49** | 6.63E-06 |
| *YDL085C-A / YDL085C-A* | **0.49** | 1.59E-06 |
| *VMR1 / YHL035C* | **0.49** | 1.71E-06 |
| *BAR1 / YIL015W* | **0.49** | 4.01E-06 |
| *TOS4 / YLR183C* | **0.49** | 2.44E-06 |
| *RFU1 / YLR073C* | **0.49** | 5.82E-05 |
| *PRM7 / YDL038C* | **0.49** | 0.0008 |
| *COT1 / YOR316C* | **0.49** | 8.77E-07 |
| *SCW10 / YMR305C* | **0.49** | 0.0002 |
| *POL1 / YNL102W* | **0.49** | 1.22E-07 |
| *TOM20 / YGR082W* | **0.49** | 2.62E-06 |
| *ELO2 / YCR034W* | **0.48** | 0.0002 |
| *ENT4 / YLL038C* | **0.48** | 5.20E-05 |
| *HMS2 / YJR147W* | **0.48** | 3.01E-06 |
| *YDR524W-C / YDR524W-A* | **0.48** | 0.0162 |
| *YBL081W / YBL081W* | **0.48** | 0.0002 |
| *RSR1 / YGR152C* | **0.48** | 4.39E-07 |
| *ICL1 / YER065C* | **0.48** | 0.0008 |
| *CLB6 / YGR109C* | **0.48** | 0.0002 |
| *ARO8 / YGL202W* | **0.48** | 8.45E-05 |
| *ASN1 / YPR145W* | **0.48** | 1.33E-05 |
| *HIP1 / YGR191W* | **0.48** | 3.42E-06 |
| *GUA1 / YMR217W* | **0.47** | 7.68E-07 |
| *ERD1 / YDR414C* | **0.47** | 2.48E-06 |
| *RSM27 / YGR215W* | **0.47** | 1.47E-06 |
| *AIM24 / YJR080C* | **0.47** | 0.0008 |
| *YBR238C / YBR238C* | **0.47** | 0.0007 |
| *MTO1 / YGL236C* | **0.47** | 1.09E-07 |

| | | |
|---|---|---|
| *HNM1 / YGL077C* | **0.47** | 0.0014 |
| *MRPS35 / YGR165W* | **0.47** | 5.47E-06 |
| *IRC7 / YFR055W* | **0.47** | 2.78E-05 |
| *ACH1 / YBL015W* | **0.47** | 1.57E-05 |
| *ISU2 / YOR226C* | **0.47** | 1.63E-07 |
| *FUM1 / YPL262W* | **0.47** | 0.0221 |
| *YGR035W-A / YGR035W-A* | **0.47** | 0.0003 |
| *CWP1 / YKL096W* | **0.46** | 0.0028 |
| *TEF4 / YKL081W* | **0.46** | 1.39E-07 |
| *YPL199C / YPL199C* | **0.46** | 5.84E-07 |
| *YBL028C / YBL028C* | **0.46** | 0.0006 |
| *MSW1 / YDR268W* | **0.46** | 0.0031 |
| *PBI1 / YPL272C* | **0.46** | 1.46E-07 |
| *SRP40 / YKR092C* | **0.46** | 3.54E-06 |
| *ADH1 / YOL086C* | **0.46** | 0.0064 |
| *RPS10A / YOR293W* | **0.46** | 6.57E-06 |
| *MNP1 / YGL068W* | **0.46** | 1.46E-05 |
| *CSI2 / YOL007C* | **0.46** | 1.93E-07 |
| *GND1 / YHR183W* | **0.46** | 0.0037 |
| *YLR264C-A / YLR264C-A* | **0.46** | 0.0037 |
| *BNA1 / YJR025C* | **0.46** | 0.0032 |
| *MRPS8 / YMR158W* | **0.45** | 5.70E-07 |
| *MIR1 / YJR077C* | **0.45** | 5.58E-05 |
| *GIC2 / YDR309C* | **0.45** | 7.09E-08 |
| *RPL18A; RPL18B / YNL301C* | **0.45** | 1.03E-05 |
| *MRPL4 / YLR439W* | **0.44** | 1.97E-06 |
| *PAM18 / YLR008C* | **0.44** | 7.77E-08 |
| *PRY2 / YKR013W* | **0.44** | 1.41E-07 |
| *MIC60 / YKR016W* | **0.44** | 1.05E-07 |
| *ATP20 / YPR020W* | **0.44** | 4.24E-08 |
| *MIS1 / YBR084W* | **0.44** | 1.67E-07 |
| *YLR042C / YLR042C* | **0.44** | 9.48E-06 |
| *DBP2 / YNL112W* | **0.44** | 0.0014 |
| *NOP56 / YLR197W* | **0.44** | 9.44E-06 |
| *CLU1 / YMR012W* | **0.44** | 1.79E-07 |
| *YAH1 / YPL252C* | **0.43** | 1.67E-07 |
| *CYS3 / YAL012W* | **0.43** | 5.20E-05 |
| *INO1 / YJL153C* | **0.43** | 0.0014 |
| *FUR4 / YBR021W* | **0.43** | 5.27E-06 |
| *TOM6 / YOR045W* | **0.43** | 9.31E-08 |
| *YHM2 / YMR241W* | **0.43** | 5.94E-09 |
| *RPM2 / YML091C* | **0.43** | 4.71E-05 |
| *YLR342W-A / YLR342W-A* | **0.42** | 1.16E-05 |
| *MRPL7 / YDR237W* | **0.42** | 6.39E-06 |
| *YLR307C-A / YLR307C-A* | **0.42** | 8.43E-07 |

| | | |
|---|---|---|
| *RPL29 / YFR032C-A* | **0.42** | 0.0001 |
| *ECM13 / YBL043W* | **0.42** | 0.0002 |
| *NUC1 / YJL208C* | **0.42** | 2.67E-08 |
| *ACS2 / YLR153C* | **0.42** | 2.34E-05 |
| *BSC1 / YDL037C* | **0.42** | 0.0007 |
| *FRE3 / YOR381W* | **0.41** | 1.24E-06 |
| *RPS8A; RPS8B / YBL072C* | **0.40** | 1.33E-06 |
| *GAS1 / YMR307W* | **0.40** | 9.39E-05 |
| *SAH1 / YER043C* | **0.40** | 9.07E-05 |
| *IMD2 / YAR073W* | **0.40** | 0.0048 |
| *RPL9A / YGL147C* | **0.39** | 1.88E-05 |
| *MET13 / YGL125W* | **0.39** | 0.0100 |
| *SRL1 / YOR247W* | **0.39** | 0.0064 |
| *OAC1 / YKL120W* | **0.38** | 1.78E-08 |
| *SIT1 / YEL065W* | **0.38** | 2.40E-07 |
| *AIM33 / YML087C* | **0.38** | 6.01E-08 |
| *CLN2 / YPL256C* | **0.38** | 1.41E-07 |
| *IMD4 / YML056C* | **0.37** | 5.69E-07 |
| *YLL053C / YLL053C* | **0.37** | 5.44E-07 |
| *ARN2 / YHL047C* | **0.37** | 1.41E-07 |
| *snR73 / YMR013W-A* | **0.36** | 3.87E-06 |
| *LEU9 / YOR108W* | **0.36** | 1.19E-08 |
| *AAH1 / YNL141W* | **0.36** | 1.73E-05 |
| *YJL047C-A / YJL047C-A* | **0.36** | 4.60E-08 |
| *YMR230W-A / YMR230W-A* | **0.36** | 0.0024 |
| *TOS6 / YNL300W* | **0.36** | 2.67E-08 |
| *AQR1 / YNL065W* | **0.35** | 2.38E-05 |
| *RNR1 / YER070W* | **0.35** | 1.28E-07 |
| *TUF1 / YOR187W* | **0.35** | 8.97E-07 |
| *QDR2 / YIL121W* | **0.35** | 2.90E-05 |
| *CYC1 / YJR048W* | **0.35** | 7.10E-08 |
| *RRT5 / YFR032C* | **0.34** | 5.51E-05 |
| *COX17 / YLL009C* | **0.34** | 2.61E-08 |
| *SPO19 / YPL130W* | **0.34** | 0.0001 |
| *MCD1 / YDL003W* | **0.34** | 2.66E-09 |
| *YHB1 / YGR234W* | **0.34** | 1.76E-08 |
| *FRE5 / YOR384W* | **0.34** | 3.74E-05 |
| *UPS2 / YLR168C* | **0.33** | 2.40E-07 |
| *GGC1 / YDL198C* | **0.32** | 6.73E-08 |
| *FIT3 / YOR383C* | **0.32** | 0.0174 |
| *IDH1 / YNL037C* | **0.31** | 2.40E-05 |
| *CIN2 / YPL241C* | **0.31** | 0.0001 |
| *PCK1 / YKR097W* | **0.30** | 4.90E-09 |
| *SAM2 / YDR502C* | **0.29** | 2.92E-08 |
| *UTR2 / YEL040W* | **0.29** | 1.80E-08 |

| | | |
|---|---|---|
| *IDH2 / YOR136W* | **0.28** | 3.11E-05 |
| *YNL042W-B; YOL013W-A; YOR072W-B / YOR072W-B* | **0.28** | 3.97E-08 |
| *YLR179C / YLR179C* | **0.28** | 5.99E-07 |
| *ATF2 / YGR177C* | **0.28** | 3.85E-07 |
| *YKL068W-A / YKL068W-A* | **0.27** | 1.91E-07 |
| *ERG4 / YGL012W* | **0.27** | 2.67E-05 |
| *FTR1 / YER145C* | **0.26** | 8.41E-09 |
| *HTA2 / YBL003C* | **0.26** | 8.48E-05 |
| *YGR035C / YGR035C* | **0.26** | 4.39E-07 |
| *TIS11 / YLR136C* | **0.25** | 0.0001 |
| *HTB2 / YBL002W* | **0.25** | 6.17E-05 |
| *IZH4 / YOL101C* | **0.23** | 3.97E-08 |
| *SAM1 / YLR180W* | **0.19** | 4.86E-10 |
| *HO / YDL227C* | **0.14** | 6.28E-09 |
| *FET3 / YMR058W* | **0.10** | 7.61E-07 |
| *FIT2 / YOR382W* | **0.10** | 0.0007 |

**Table S2. Yeast genes differentially expressed after IRAK2 expression.**

| Gene name | Fold change | FDR |
|---|---|---|
| **UPREGULATED** | | |
| *SSA4 / YER103W* | **4.81** | 1.48E-08 |
| *BTN2 / YGR142W* | **3.48** | 2.30E-05 |
| *CUR1 / YPR158W* | **2.95** | 5.88E-06 |
| *HXT11 / YOL156W* | **2.43** | 0.0012 |
| *HXT9 / YJL219W* | **2.22** | 0.0168 |
| *YDR222W / YDR222W* | **2.13** | 0.0107 |
| *HSP82 / YPL240C* | **2.1** | 0.0002 |
| *MBF1 / YOR298C-A* | **2.05** | 0.0001 |
| *YNL146C-A / YNL146C-A* | **2.01** | 0.0017 |
| *REG2 / YBR050C* | **2** | 0.0001 |
| **DOWNREGULATED** | | |
| *BAG7 / YOR134W* | **0.48** | 0.0003 |
| *TIS11 / YLR136C* | **0.48** | 0.0467 |
| *YBL005W-A / YBL005W-A* | **0.47** | 0.0084 |
| *GTO3 / YMR251W* | **0.46** | 0.0016 |
| *BSC1 / YDL037C* | **0.40** | 0.0071 |
| *PHD1 / YKL043W* | **0.39** | 0.0001 |

**Table S3. Yeast genes differentially expressed after IRAK4 expression.**

| Gene name | Fold change | FDR |
|---|---|---|
| **UPREGULATED** | | |
| *GRE1 / YPL223C* | **12.29** | 0.0002 |
| *CTT1 / YGR088W* | **12.1** | 4.03E-10 |
| *FMP48 / YGR052W* | **10.95** | 4.70E-10 |
| *TKL2 / YBR117C* | **10.77** | 1.16E-08 |
| *RTC3 / YHR087W* | **8.97** | 3.17E-08 |
| *GND2 / YGR256W* | **8.76** | 1.82E-06 |
| *ALD3 / YMR169C* | **8.73** | 4.80E-07 |
| *RTN2 / YDL204W* | **8.4** | 5.05E-08 |
| *YNR014W* | **7.96** | 2.30E-05 |
| *FMP45 / YDL222C* | **7.57** | 2.62E-09 |
| *YKL107W / YKL107W* | **7.42** | 8.58E-05 |
| *PAI3 / YMR174C* | **6.72** | 2.08E-05 |
| *PHM7 / YOL084W* | **6.4** | 9.49E-07 |
| *GAD1 / YMR250W* | **5.99** | 2.68E-08 |
| *SOL4 / YGR248W* | **5.83** | 2.42E-09 |
| *SIP18 / YMR175W* | **5.83** | 0.0005 |
| *HPF1/ YIL169C* | **5.68** | 6.49E-06 |
| *YDL218W / YDL218W* | **5.4** | 1.77E-05 |
| *MSC1 / YML128C* | **5.22** | 6.24E-10 |
| *HBT1 / YDL223C* | **5.06** | 2.06E-07 |
| *TSA2 / YDR453C* | **4.99** | 0.0041 |
| *NQM1 / YGR043C* | **4.85** | 3.27E-08 |
| *SPI1 / YER150W* | **4.85** | 2.94E-08 |
| *HPF1 / YOL155C* | **4.81** | 2.84E-05 |
| *PMA2 / YPL036W* | **4.8** | 8.68E-06 |
| *HSP31 / YDR533C* | **4.7** | 0.0005 |
| *CHA1 / YCL064C* | **4.62** | 0.0002 |
| *TFS1 / YLR178C* | **4.35** | 1.62E-09 |
| *SSA3 / YBL075C* | **4.29** | 6.07E-05 |
| *HSP26 / YBR072W* | **4.2** | 2.96E-06 |
| *YGP1 / YNL160W* | **4.19** | 9.43E-10 |
| *YJL045W / YJL045W* | **4.11** | 0.0034 |
| *FMP16 / YDR070C* | **4.02** | 1.38E-06 |
| *YJR096W / YJR096W* | **4** | 4.70E-08 |
| *GIP2 / YER054C* | **3.99** | 2.66E-07 |
| *YJL144W / YJL144W* | **3.96** | 1.09E-06 |
| *DCS2 / YOR173W* | **3.89** | 3.13E-09 |
| *SHH3 / YMR118C* | **3.77** | 0.0371 |
| *YNL194C / YNL194C* | **3.73** | 2.78E-06 |
| *NRG1 / YDR043C* | **3.71** | 1.62E-09 |

| | | |
|---|---|---|
| *RPI1 / YIL119C* | **3.67** | 9.32E-09 |
| *SHH4 / YLR164W* | **3.58** | 4.67E-05 |
| *VPS73 / YGL104C* | **3.55** | 9.32E-09 |
| *YDR034W-B* | **3.51** | 0.0003 |
| *YGR066C* | **3.46** | 0.0031 |
| *YOR186W* | **3.44** | 0.0009 |
| *CIN5 / YOR028C* | **3.42** | 2.67E-07 |
| *PES4 / YFR023W* | **3.35** | 4.80E-07 |
| *YJL163C / YJL163C* | **3.34** | 1.70E-07 |
| *PHM8 / YER037W* | **3.31** | 0.0003 |
| *COX5B / YIL111W* | **3.3** | 8.67E-08 |
| *FUN19 / YAL034C* | **3.27** | 3.27E-08 |
| *BDH2 / YAL061W* | **3.23** | 8.83E-08 |
| *HXT5 / YHR096C* | **3.16** | 1.18E-06 |
| *TSL1 / YML100W* | **3.14** | 2.83E-07 |
| *YGL258W-A* | **3.12** | 0.0001 |
| *YPT53 / YNL093W* | **3.11** | 1.67E-06 |
| *TDH1 / YJL052W* | **3.05** | 0.0002 |
| *YNL195C / YNL195C* | **3.05** | 1.99E-07 |
| *YJL016W / YJL016W* | **3.05** | 9.32E-09 |
| *YPL247C / YPL247C* | **3.05** | 3.14E-08 |
| *SYM1 / YLR251W* | **3.04** | 9.32E-09 |
| *GPG1 / YGL121C* | **3.03** | 1.33E-06 |
| *BAG7 / YOR134W* | **3.03** | 1.70E-07 |
| *FRT2 / YAL028W* | **2.99** | 9.32E-09 |
| *SPS100 / YHR139C* | **2.97** | 0.0041 |
| *SDP1 / YIL113W* | **2.96** | 2.02E-06 |
| *YMR090W* | **2.92** | 0.0004 |
| *MPO1 / YGL010W* | **2.9** | 9.32E-09 |
| *HSP42 / YDR171W* | **2.88** | 5.05E-08 |
| *MCH2 / YKL221W* | **2.86** | 2.48E-05 |
| *WSC4 / YHL028W* | **2.86** | 3.26E-07 |
| *GPD1 / YDL022W* | **2.84** | 9.32E-09 |
| *TOS8 / YGL096W* | **2.84** | 4.78E-06 |
| *AQY1 / YPR192W* | **2.84** | 0.0102 |
| *YER039C-A* | **2.82** | 3.07E-06 |
| *ATG29 / YPL166W* | **2.81** | 0.0004 |
| *OCH1 / YGL038C* | **2.8** | 2.94E-08 |
| *ERR1; ERR2; ERR3 / YMR323W* | **2.76** | 0.0017 |
| *ROM1 / YGR070W* | **2.73** | 8.93E-07 |
| *SPG1 / YGR236C* | **2.71** | 2.56E-05 |
| *YJR115W / YJR115W* | **2.7** | 3.76E-06 |
| *XBP1 / YIL101C* | **2.7** | 4.52E-06 |
| *OM45 / YIL136W* | **2.7** | 8.48E-07 |
| *SHC1 / YER096W* | **2.7** | 0.0009 |

| | | |
|---|---|---|
| *HSP30 / YCR021C* | **2.68** | 0.0008 |
| *PRY2 / YKR013W* | **2.67** | 6.25E-08 |
| *TPS2 / YDR074W* | **2.66** | 1.37E-06 |
| *RCK1 / YGL158W* | **2.65** | 0.0322 |
| *GRE2 / YOL151W* | **2.63** | 0.0001 |
| *PNS1 / YOR161C* | **2.62** | 1.05E-05 |
| *HSP32; HSP33; SNO4 / YMR322C* | **2.58** | 2.96E-07 |
| *SDS24 / YBR214W* | **2.58** | 1.09E-06 |
| *PIR3 / YKL163W* | **2.56** | 0.0004 |
| *YPL014W / YPL014W* | **2.55** | 0.0001 |
| *ECM34 / YHL043W* | **2.54** | 1.00E-05 |
| *PIC2 / YER053C* | **2.53** | 7.77E-08 |
| *CRG1 / YHR209W* | **2.53** | 5.83E-07 |
| *PHO3 / YBR092C* | **2.53** | 0.0443 |
| *GGA1 / YDR358W* | **2.52** | 4.29E-07 |
| *UIP4 / YPL186C* | **2.51** | 8.12E-07 |
| *UGP1 / YKL035W* | **2.5** | 2.14E-06 |
| *TPS1 / YBR126C* | **2.5** | 6.38E-07 |
| *PCL1 / YNL289W* | **2.5** | 3.92E-08 |
| *YKL151C / YKL151C* | **2.48** | 5.44E-05 |
| *GPI2 / YPL076W* | **2.47** | 6.77E-06 |
| *ENO1 / YGR254W* | **2.46** | 9.51E-06 |
| *YHL012W / YHL012W* | **2.45** | 2.05E-05 |
| *IMP2' / YIL154C* | **2.44** | 2.18E-05 |
| *YLR031W / YLR031W* | **2.42** | 0.0004 |
| *CSI2 / YOL007C* | **2.42** | 7.22E-07 |
| *YGR201C / YGR201C* | **2.42** | 0.0017 |
| *RCN2 / YOR220W* | **2.42** | 4.25E-07 |
| *PHO5 / YBR093C* | **2.4** | 4.57E-06 |
| *YDL124W / YDL124W* | **2.4** | 1.35E-08 |
| *CLN2 / YPL256C* | **2.38** | 1.55E-06 |
| *SAF1 / YBR280C* | **2.38** | 5.45E-07 |
| *APE1 / YKL103C* | **2.36** | 1.21E-07 |
| *YRO2 / YBR054W* | **2.36** | 5.28E-06 |
| *GSY2 / YLR258W* | **2.35** | 1.33E-06 |
| *DBP1 / YPL119C* | **2.33** | 5.97E-06 |
| *YIL077C / YIL077C* | **2.32** | 1.09E-07 |
| *YIL169C / YIL169C* | **2.32** | 0.0002 |
| *SVS1 / YPL163C* | **2.31** | 9.49E-07 |
| *CRS5 / YOR031W* | **2.3** | 0.0027 |
| *CLB1 / YGR108W* | **2.3** | 4.29E-07 |
| *YHL042W / YHL042W* | **2.3** | 1.69E-05 |
| *URA10 / YMR271C* | **2.29** | 4.05E-05 |
| *ATG8 / YBL078C* | **2.28** | 1.35E-05 |
| *RME1 / YGR044C* | **2.27** | 5.62E-05 |

| | | |
|---|---|---|
| *SRL3 / YKR091W* | **2.26** | 3.06E-05 |
| *YLR257W / YLR257W* | **2.24** | 1.11E-05 |
| *TOS6 / YNL300W* | **2.24** | 8.08E-07 |
| *NCA3 / YJL116C* | **2.23** | 0.0096 |
| *GPM2 / YDL021W* | **2.23** | 4.01E-06 |
| *PLM2 / YDR501W* | **2.22** | 5.08E-06 |
| *FYV10 / YIL097W* | **2.21** | 5.11E-06 |
| *GTT1 / YIR038C* | **2.21** | 9.51E-06 |
| *HVG1 / YER039C* | **2.21** | 3.92E-06 |
| *AMS1 / YGL156W* | **2.2** | 5.52E-07 |
| *PCL2 / YDL127W* | **2.2** | 5.97E-06 |
| *DIA3 / YDL024C* | **2.19** | 0.0008 |
| *YNL200C / YNL200C* | **2.19** | 1.88E-05 |
| *HMS1 / YOR032C* | **2.18** | 0.0014 |
| *YAK1 / YJL141C* | **2.17** | 5.28E-06 |
| *YHR097C / YHR097C* | **2.17** | 4.94E-07 |
| *PGM2 / YMR105C* | **2.17** | 4.29E-07 |
| *USV1 / YPL230W* | **2.16** | 1.12E-05 |
| *NTH1 / YDR001C* | **2.16** | 8.68E-06 |
| *ATG9 / YDL149W* | **2.15** | 5.56E-06 |
| *YGL036W / YGL036W* | **2.14** | 4.05E-05 |
| *DDR2 / YOL052C-A* | **2.13** | 7.69E-06 |
| *SSA4 / YER103W* | **2.13** | 2.33E-06 |
| *GSP2 / YOR185C* | **2.13** | 3.22E-05 |
| *TPK2 / YPL203W* | **2.12** | 7.69E-06 |
| *MGA1 / YGR249W* | **2.12** | 8.01E-05 |
| *GPX1 / YKL026C* | **2.12** | 3.37E-06 |
| *TPK1 / YJL164C* | **2.11** | 2.93E-07 |
| *CYC7 / YEL039C* | **2.1** | 4.14E-05 |
| *PDE1 / YGL248W* | **2.1** | 6.30E-06 |
| *PRB1 / YEL060C* | **2.09** | 2.28E-07 |
| *MUP3 / YHL036W* | **2.09** | 1.33E-06 |
| *YNR064C / YNR064C* | **2.09** | 2.04E-05 |
| *BXI1 / YNL305C* | **2.09** | 7.22E-07 |
| *MIT1 / YEL007W* | **2.08** | 0.0002 |
| *LDS2 / YOL047C* | **2.07** | 0.0001 |
| *DAK2 / YFL053W* | **2.07** | 0.0360 |
| *FMP33 / YJL161W* | **2.06** | 3.60E-05 |
| *IKS1 / YJL057C* | **2.06** | 9.49E-06 |
| *SSE2 / YBR169C* | **2.06** | 2.31E-05 |
| *YIL029C / YIL029C* | **2.04** | 0.0002 |
| *ATG19 / YOL082W* | **2.04** | 4.17E-06 |
| *IRC15 / YPL017C* | **2.04** | 5.96E-05 |
| *IGD1 / YFR017C* | **2.04** | 3.76E-06 |
| *RNY1 / YPL123C* | **2.03** | 1.99E-06 |

| | | |
|---|---|---|
| *DUN1 / YDL101C* | **2.02** | 1.08E-06 |
| *PST1 / YDR055W* | **2.01** | 3.22E-05 |
| *EDC2 / YER035W* | **2** | 1.71E-05 |
| *GLO4 / YOR040W* | **2** | 0.0429 |
| *YMR196W* | **2** | 9.64E-05 |
| **DOWNREGULATED** | | |
| *ENB1 / YOL158C* | **0.50** | 8.51E-05 |
| *HXK2 / YGL253W* | **0.50** | 4.19E-05 |
| *AAT1 / YKL106W* | **0.50** | 0.0029 |
| *ICL1 / YER065C* | **0.50** | 0.0002 |
| *YER138W-A; YOR192C-C* | **0.50** | 0.0132 |
| *NRG2 / YBR066C* | **0.50** | 0.0002 |
| *GTO3 / YMR251W* | **0.49** | 0.0002 |
| *BAT2 / YJR148W* | **0.49** | 0.0001 |
| *IDP3 / YNL009W* | **0.49** | 0.0010 |
| *ICS3 / YJL077C* | **0.49** | 0.0143 |
| *MCM10 / YIL150C* | **0.49** | 0.0198 |
| *CTF13 / YMR094W* | **0.49** | 0.0039 |
| *IMD2 / YAR073W* | **0.49** | 0.0060 |
| *BMT6 / YLR063W* | **0.48** | 2.40E-05 |
| *SSA2 / YLL024C* | **0.48** | 1.70E-07 |
| *PEX21 / YGR239C* | **0.48** | 2.70E-06 |
| *EEB1 / YPL095C* | **0.48** | 0.0031 |
| *TAH1 / YCR060W* | **0.48** | 6.52E-06 |
| *AAP1 / YHR047C* | **0.48** | 1.68E-05 |
| *PTR2 / YKR093W* | **0.48** | 0.0124 |
| *RPL41A; RPL41B / YDL184C* | **0.47** | 0.0007 |
| *ATO3 / YDR384C* | **0.47** | 1.59E-05 |
| *REE1 / YJL217W* | **0.47** | 2.87E-06 |
| *ARO9 / YHR137W* | **0.47** | 3.05E-05 |
| *DOG2 / YHR043C* | **0.47** | 1.12E-05 |
| *DBR1 / YKL149C* | **0.46** | 7.30E-06 |
| *YLR363W-A* | **0.46** | 0.0193 |
| *FTR1 / YER145C* | **0.46** | 3.76E-06 |
| *MCH5 / YOR306C* | **0.46** | 1.39E-05 |
| *PRM4 / YPL156C* | **0.46** | 4.80E-07 |
| *SAM4 / YPL273W* | **0.46** | 9.70E-08 |
| *PIR5 / YJL160C* | **0.45** | 0.0001 |
| *CYB2 / YML054C* | **0.45** | 0.0375 |
| *SRL4 / YPL033C* | **0.45** | 0.0210 |
| *SSP1 / YHR184W* | **0.45** | 0.0007 |
| *37226 / YLR348C* | **0.44** | 0.0029 |
| *MRK1 / YDL079C* | **0.44** | 1.33E-06 |
| *SAM4 / YMR321C* | **0.44** | 1.02E-07 |
| *NAT4 / YMR069W* | **0.44** | 0.0033 |

| | | |
|---|---|---|
| *GUT1 / YHL032C* | **0.43** | 0.0004 |
| *YBL005W-A* | **0.43** | 0.0002 |
| *MMP1 / YLL061W* | **0.43** | 6.52E-06 |
| *UBC11 / YOR339C* | **0.43** | 0.0014 |
| *ARN2 / YHL047C* | **0.42** | 1.66E-06 |
| *MAL12; MAL32 / YBR299W* | **0.42** | 1.77E-07 |
| *AAH1 / YNL141W* | **0.42** | 0.0004 |
| *YJL047C-A / YJL047C-A* | **0.42** | 4.80E-07 |
| *FAA4 / YMR246W* | **0.41** | 4.78E-06 |
| *REG2 / YBR050C* | **0.41** | 5.87E-08 |
| *DCI1 / YOR180C* | **0.41** | 0.0112 |
| *DBP2 / YNL112W* | **0.41** | 0.0232 |
| *IMD2 / YHR216W* | **0.40** | 0.0016 |
| *OYE2 / YHR179W* | **0.40** | 1.22E-05 |
| *FIT2 / YOR382W* | **0.39** | 0.0469 |
| *FMP23 / YBR047W* | **0.39** | 8.73E-07 |
| *YGR035C / YGR035C* | **0.39** | 6.13E-06 |
| *TIS11 / YLR136C* | **0.39** | 0.0011 |
| *ALP1 / YNL270C* | **0.38** | 0.0178 |
| *FRE5 / YOR384W* | **0.38** | 3.03E-05 |
| *YBL028C / YBL028C* | **0.38** | 0.0011 |
| *FRE3 / YOR381W* | **0.38** | 9.39E-07 |
| *YMC2 / YBR104W* | **0.37** | 3.67E-08 |
| *snR73 / YMR013W-A* | **0.37** | 2.93E-05 |
| *SIT1 / YEL065W* | **0.36** | 1.21E-07 |
| *RRT5 / YFR032C* | **0.36** | 0.0002 |
| *YML131W / YML131W* | **0.36** | 0.0001 |
| *CAT2 / YML042W* | **0.36** | 3.21E-05 |
| *SPO19 / YPL130W* | **0.36** | 0.0003 |
| *FOX2 / YKR009C* | **0.36** | 1.71E-05 |
| *TAT1 / YBR069C* | **0.36** | 1.10E-06 |
| *YGR067C / YGR067C* | **0.36** | 0.0013 |
| *YDL085C-A / YDL085C-A* | **0.36** | 5.87E-08 |
| *YLR307C-A / YLR307C-A* | **0.36** | 3.06E-07 |
| *MLS1 / YNL117W* | **0.35** | 1.16E-08 |
| *RAD55 / YDR076W* | **0.35** | 9.51E-06 |
| *MAL33 / YBR297W* | **0.35** | 4.29E-07 |
| *MIG2 / YGL209W* | **0.34** | 0.0019 |
| *YKL068W-A* | **0.34** | 5.56E-06 |
| *SPS19 / YNL202W* | **0.34** | 8.68E-06 |
| *CAR1 / YPL111W* | **0.34** | 7.69E-06 |
| *ECI1 / YLR284C* | **0.33** | 1.14E-05 |
| *YMR230W-A* | **0.33** | 0.0241 |
| *YKR075C / YKR075C* | **0.32** | 8.72E-06 |
| *CRC1 / YOR100C* | **0.32** | 0.0004 |

| | | |
|---|---|---|
| *HBN1 / YCL026C-B* | **0.32** | 0.0001 |
| *INO1 / YJL153C* | **0.32** | 0.0001 |
| *EFG1 / YGR272C* | **0.31** | 0.0001 |
| *IMA5 / YJL216C* | **0.31** | 1.58E-05 |
| *CSM4 / YPL200W* | **0.28** | 4.87E-06 |
| *HXT4 / YHR092C* | **0.27** | 3.02E-08 |
| *FAT3 / YKL187C* | **0.27** | 3.67E-08 |
| *SUC2 / YIL162W* | **0.27** | 6.17E-09 |
| *PXA1 / YPL147W* | **0.26** | 3.67E-08 |
| *PRM7 / YDL038C* | **0.26** | 3.73E-06 |
| *PRM7 / YDL039C* | **0.23** | 1.39E-05 |
| *PDH1 / YPR002W* | **0.23** | 1.22E-05 |
| *PHO84 / YML123C* | **0.23** | 0.0164 |
| *BSC1 / YDL037C* | **0.23** | 5.84E-06 |
| *CIT3 / YPR001W* | **0.22** | 2.41E-05 |
| *LPX1 / YOR084W* | **0.22** | 0.0002 |
| *ACS1 / YAL054C* | **0.22** | 3.84E-08 |
| *FAA2 / YER015W* | **0.22** | 1.99E-06 |
| *RGI2 / YIL057C* | **0.20** | 6.61E-07 |
| *POX1 / YGL205W* | **0.20** | 2.02E-06 |
| *CTA1 / YDR256C* | **0.19** | 1.72E-07 |
| *YOR072W-B* | **0.19** | 3.41E-09 |
| *TIP1 / YBR067C* | **0.18** | 9.43E-10 |
| *YIG1 / YPL201C* | **0.18** | 2.23E-08 |
| *FIT3 / YOR383C* | **0.18** | 4.35E-05 |
| *ADY2 / YCR010C* | **0.16** | 7.79E-08 |
| *ICL2 / YPR006C* | **0.15** | 3.57E-05 |
| *FDH1 / YOR388C* | **0.14** | 3.71E-05 |
| *ADH2 / YMR303C* | **0.12** | 4.03E-10 |
| *HXT2 / YMR011W* | **0.12** | 9.86E-07 |
| *SUL1 / YBR294W* | **0.11** | 4.70E-08 |

**Table S4. Yeast genes differentially expressed after IRAK4 (KD) expression.**

| Gene name | Fold change | FDR |
|---|---|---|
| **UPREGULATED** | | |
| *YOR387C / YOR387C* | **15.76** | 8.86E-07 |
| *VEL1 / YGL258W* | **8.58** | 4.38E-05 |
| *ZPS1 / YOL154W* | **4.52** | 9.25E-07 |
| *SPI1 / YER150W* | **3.19** | 6.05E-06 |
| *CHA1 / YCL064C* | **3.01** | 0.0085 |
| *HSP26 / YBR072W* | **2.74** | 0.0004 |
| *SSA3 / YBL075C* | **2.72** | 0.0098 |
| *SSA4 / YER103W* | **2.71** | 1.34E-06 |
| *ADH4 / YGL256W* | **2.7** | 5.48E-06 |
| *BTN2 / YGR142W* | **2.61** | 3.30E-05 |
| *YER053C-A / YER053C-A* | **2.56** | 4.38E-05 |
| *HSP42 / YDR171W* | **2.38** | 5.45E-06 |
| *HSP30 / YCR021C* | **2.32** | 0.0102 |
| *YRO2 / YBR054W* | **2.31** | 0.0001 |
| *YGP1 / YNL160W* | **2.29** | 3.95E-06 |
| *SUC2 / YIL162W* | **2.27** | 1.88E-05 |
| *CUR1 / YPR158W* | **2.24** | 1.18E-05 |
| *INA1 / YLR413W* | **2.2** | 0.0003 |
| *FRE7 / YOL152W* | **2.17** | 0.0022 |
| *HXT6; HXT7 / YDR342C* | **2.14** | 0.0003 |
| *HSP82 / YPL240C* | **2.11** | 5.19E-05 |
| *TDA6 / YPR157W* | **2.08** | 1.13E-05 |
| *SAM2 / YDR502C* | **2.07** | 0.0003 |
| *CTR1 / YPR124W* | **2.06** | 0.0042 |
| *TDH1 / YJL052W* | **2.05** | 0.0259 |
| *OPI10 / YOL032W* | **2** | 9.60E-05 |
| **DOWNREGULATED** | | |
| *SNO1 / YMR095C* | **0.5** | 0.0053 |
| *ICY2 / YPL250C* | **0.5** | 0.0033 |
| *ZTA1 / YBR046C* | **0.5** | 1.55E-05 |
| *YBL028C / YBL028C* | **0.49** | 0.0297 |
| *PXA1 / YPL147W* | **0.49** | 0.0002 |
| *MAE1 / YKL029C* | **0.49** | 0.0058 |
| *RGM1 / YMR182C* | **0.49** | 0.0018 |
| *IDP3 / YNL009W* | **0.48** | 0.0053 |
| *ARO9 / YHR137W* | **0.48** | 0.0003 |
| *CIN1 / YOR349W* | **0.48** | 0.0003 |
| *ARO10 / YDR380W* | **0.48** | 0.0006 |
| *YAT1 / YAR035W* | **0.48** | 0.0037 |
| *YAR035C-A / YAR035C-A* | **0.46** | 0.0074 |

| | | |
|---|---|---|
| *ARN2 / YHL047C* | **0.46** | 6.76E-05 |
| *SPG4 / YMR107W* | **0.46** | 0.0011 |
| *YGR035C / YGR035C* | **0.46** | 0.0020 |
| *YOR072W-B / YOR072W-B* | **0.45** | 0.0003 |
| *CWC25 / YNL245C* | **0.45** | 0.0011 |
| *SSU1 / YPL092W* | **0.45** | 7.03E-05 |
| *YLR307C-A / YLR307C-A* | **0.45** | 6.05E-06 |
| *HXT4 / YHR092C* | **0.44** | 2.08E-05 |
| *CAT2 / YML042W* | **0.44** | 0.0009 |
| *ECI1 / YLR284C* | **0.44** | 0.0005 |
| *MMS21 / YEL019C* | **0.44** | 0.0070 |
| *DCI1 / YOR180C* | **0.44** | 0.0282 |
| *YKR075C / YKR075C* | **0.44** | 0.0003 |
| *FMP23 / YBR047W* | **0.43** | 1.18E-05 |
| *YDR034W-B / YDR034W-B* | **0.43** | 0.0465 |
| *SPO19 / YPL130W* | **0.43** | 0.0065 |
| *FIT3 / YOR383C* | **0.42** | 0.0159 |
| *ENT4 / YLL038C* | **0.42** | 8.83E-05 |
| *NDE2 / YDL085W* | **0.42** | 9.60E-05 |
| *CRF1 / YDR223W* | **0.41** | 0.0025 |
| *MCM10 / YIL150C* | **0.41** | 0.0108 |
| *PUT4 / YOR348C* | **0.41** | 0.0024 |
| *FAT3 / YKL187C* | **0.40** | 1.10E-05 |
| *FOX2 / YKR009C* | **0.40** | 0.0002 |
| *HXT3 / YDR345C* | **0.40** | 6.52E-05 |
| *MDH2 / YOL126C* | **0.39** | 9.25E-07 |
| *BOP2 / YLR267W* | **0.39** | 2.92E-05 |
| *CTF13 / YMR094W* | **0.38** | 0.0034 |
| *FAA2 / YER015W* | **0.38** | 0.0016 |
| *AQR1 / YNL065W* | **0.37** | 0.0007 |
| *ADH2 / YMR303C* | **0.36** | 3.95E-06 |
| *FBP1 / YLR377C* | **0.36** | 9.90E-07 |
| *PRM7 / YDL038C* | **0.33** | 0.0002 |
| *PIR5 / YJL160C* | **0.33** | 1.66E-05 |
| *PRM7 / YDL039C* | **0.33** | 0.0007 |
| *POX1 / YGL205W* | **0.32** | 0.0004 |
| *POT1 / YIL160C* | **0.31** | 0.0003 |
| *GAP1 / YKR039W* | **0.31** | 1.56E-05 |
| *CRC1 / YOR100C* | **0.31** | 0.0010 |
| *ACS1 / YAL054C* | **0.30** | 3.14E-06 |
| *CIT3 / YPR001W* | **0.30** | 0.0009 |
| *YBL005W-A / YBL005W-A* | **0.30** | 0.0002 |
| *PCK1 / YKR097W* | **0.30** | 2.49E-07 |
| *LPX1 / YOR084W* | **0.28** | 0.0056 |
| *ICL2 / YPR006C* | **0.27** | 0.0039 |

| | | |
|---|---|---|
| *CTA1 / YDR256C* | **0.26** | 1.18E-05 |
| *MLS1 / YNL117W* | **0.26** | 1.02E-08 |
| *RRT5 / YFR032C* | **0.26** | 0.0006 |
| *BSC1 / YDL037C* | **0.26** | 0.0001 |
| *HXT2 / YMR011W* | **0.21** | 6.76E-05 |
| *MIG2 / YGL209W* | **0.19** | 4.99E-05 |
| *ADY2 / YCR010C* | **0.18** | 2.09E-06 |
| *SUL1 / YBR294W* | **0.07** | 8.12E-08 |

**Table S5. Biological processes regulated by the heterologous expression of IRAK4 in *S. cerevisiae* according to PANTHER classification.**

| INDUCED | | | REPRESSED | | |
|---|---|---|---|---|---|
| **Biological Process** | **FC** | **FDR** | **Biological Process** | **FC** | **FDR** |
| Trehalose biosynthesis. | 29.05 | 2.72 $x10^{-3}$ | Propionate metabolism. | 65.86 | 7.85 $x10^{-3}$ |
| Carbohydrate catabolism. | 4.9 | 1.56 $x10^{-2}$ | Long-chain fatty acids transport. | 32.93 | 2.43 $x10^{-3}$ |
| Response to extracellular stimuli. | 4.07 | 1.64 $x10^{-2}$ | Siderophore-dependent iron import into the cell. | 32.93 | 2.38 $x10^{-3}$ |
| Generation of precursor metabolites and energy. | 3.49 | 1.32 $x10^{-2}$ | Siderophores transmembrane transport. | 32.93 | 2.33 $x10^{-3}$ |
| | | | β-oxidation of fatty acids. | 30.4 | 6.17 $x10^{-5}$ |
| | | | Sucrose catabolism | 24.7 | 4.32 $x10^{-2}$ |
| | | | Glyoxylate cycle. | 24.7 | 4.24 $x10^{-2}$ |
| | | | Carboxylic acids transmembrane transport. | 7.53 | 2.73 $x10^{-3}$ |
| FC. fold change; FDR. false-discovery rate. | | | | | |

**Table S6. Biological processes regulated by the heterologous expression of IRAK4(KD) in *S. cerevisiae* according to PANTHER classification.**

| REPRESSED | | |
|---|---|---|
| **Biological Process** | **FC** | **FDR** |
| β-oxidation of fatty acids. | 61.56 | 7.53 $x10^{-8}$ |
| Monocarboxylic acids transport. | 16.33 | 3.13 $x10^{-2}$ |
| Tricarboxylic acid cycle (TCA). | 14.75 | 3.81 $x10^{-2}$ |
| Transmembrane transport. | 3.12 | 3.9 $x10^{-2}$ |
| FC. fold change; FDR. false-discovery rate. | | |

**Table S7. Biological processes regulated by the heterologous expression of IRAK1 in *S. cerevisiae* according to PANTHER classification.**

| INDUCED | | | REPRESSED | | |
|---|---|---|---|---|---|
| **Biological Process** | **FC** | **FDR** | **Biological Process** | **FC** | **FDR** |
| β-oxidation of fatty acids. | 11.54 | 5.1 $x10^{-4}$ | One-carbon metabolism. | 12.47 | 2.05 $x10^{-2}$ |

| | | | | | |
|---|---|---|---|---|---|
| Glycogen biosynthesis. | 11.54 | 4.98 x10$^{-4}$ | Iron ion transmembrane transport. | 10.13 | 5.69 x10$^{-3}$ |
| Trehalose metabolism. | 8.53 | 4.21 x10$^{-2}$ | Intracellular iron ion homeostasis. | 7.62 | 4.4 x10$^{-3}$ |
| Aldehyde catabolism. | 8.53 | 4.17 x10$^{-2}$ | Methionine metabolism. | 7.48 | 4.49 x10$^{-2}$ |
| Catabolism of oligosaccharides. | 8.21 | 6.48 x10$^{-3}$ | Sulfur compound biosynthesis. | 4.99 | 4.92 x10$^{-2}$ |
| Glucose 6-phosphate metabolism. | 7.15 | 4.65 x10$^{-3}$ | Purine compound biosynthesis | 4.44 | 4.78 x10$^{-2}$ |
| Late nucleophagy | 6.62 | 3.78 x10$^{-2}$ | Amino acid biosynthesis. | 2.52 | 4..77 x10$^{-2}$ |
| NADP metabolism. | 5.71 | 3.06 x10$^{-2}$ | Translation. | 2.55 | 1.91 x10$^{-2}$ |
| Hexoses transmembrane transport. | 5.47 | 3.7 x10$^{-2}$ | | | |
| Cellular oxidative detoxification. | 5.28 | 9.89 x10$^{-3}$ | | | |
| Oxidative stress response. | 2.65 | 4.89 x10$^{-2}$ | | | |

FC. fold change; FDR. false-discovery rate;
NADP. nicotinamide adenine dinucleotide phosphate.

**Table S8. Transcription factors involved in the regulation of differentially expressed genes found in IRAK expression datasets in S. cerevisiae.**

| Transcription factor | % genes regulated in the whole genome | % % genes regulated in the dataset | p-value | Function[a] |
|---|---|---|---|---|
| **IRAK4** | | | | |
| **Regulating induced genes** | | | | |
| Yap6 | 6.67% | 36.31% | 0 | Regulation of the expression of genes involved in carbohydrate metabolism. |
| Spt23 | 5.85% | 59.78% | 0 | Regulation of *OLE1* transcription. which is required for the synthesis of monounsaturated fatty acids and for the normal distribution of mitochondria. |

| | | | | |
|---|---|---|---|---|
| Hot1 | 16.55% | 12.85% | $1.4 \times 10^{-14}$ | Regulation of genes involved in glycerol biosynthesis in response to elevated osmolarity. |
| Sko1 | 5.95% | 35.20% | $2.18 \times 10^{-13}$ | Involved in responses to osmotic and oxidative stress. |
| Ndt80 | 4.06% | 49.16% | $3.02 \times 10^{-9}$ | Meiosis-specific transcription factor. |
| Yap5 | 4.95% | 24.58% | $5.69 \times 10^{-7}$ | Detects high iron conditions and participates in diauxic change. |
| Cad1 | 5.18% | 18.99% | $4.62 \times 10^{-6}$ | Involved in stress responses. iron metabolism. and pleiotropic drug resistance. |
| Adr1 | 4.87% | 21.23% | $5.7 \times 10^{-6}$ | Transcription of the *ADH2* gene that represses glucose. of peroxisomal protein genes. and of genes required for the utilization of ethanol. glycerol. and fatty acids. |
| **Regulating repressed genes** | | | | |
| Adr1 | 4.48% | 30.43% | $6.5 \times 10^{-10}$ | See above for genes induced by IRAK4. |
| Cat8 | 6.69% | 16.52% | $9.35 \times 10^{-9}$ | Required for gene induction under non-fermentative growth conditions. active after the diauxic shift. It binds to elements that respond to the carbon source. |
| Oaf1 | 3.74% | 32.17% | $3.21 \times 10^{-8}$ | It regulates genes involved in fatty acid β-oxidation and peroxisome organization and biogenesis. It is also involved in the diauxic shift. |
| Hap1 | 3.76% | 23.48% | $2.74 \times 10^{-6}$ | Involved in the regulation of gene expression in response to heme and oxygen levels. |

| | | | | |
|---|---|---|---|---|
| Yap6 | 3.18% | 26.96% | 1.78 $\times10^{-5}$ | See above for genes induced by IRAK4 |
| **IRAK4(KD)** | | | | |
| **Regulating induced genes** | | | | |
| Spt23 | 0.87% | 61.54% | 1.19 $\times10^{-5}$ | See above for genes induced by IRAK4. |
| **Regulating repressed genes** | | | | |
| Adr1 | 3.59% | 42.42% | 2.35 $\times10^{-12}$ | See above for genes induced by IRAK4 |
| Cat8 | 5.99% | 25.76% | 2.18 $\times10^{-11}$ | See above for genes induced by IRAK4 |
| Oaf1 | 2.83% | 42.42% | 8.04 $\times10^{-10}$ | See above for genes induced by IRAK4 |
| Rds2 | 2.62% | 22.73% | 2.73 $\times10^{-5}$ | Regulation of gluconeogenesis and the glyoxylate cycle. |
| Sip4 | 3.64% | 13.64% | 5.45 $\times10^{-5}$ | Positive regulation of gluconeogenesis. regulated by the Snf1 protein. |
| **IRAK1** | | | | |
| **Regulating induced genes** | | | | |
| Spt23 | 9.51% | 45.19% | 0 | See above for genes induced by IRAK4. |
| Yap6 | 10.87% | 27.53% | 2 $\times10^{-15}$ | See above for genes induced by IRAK4 |
| Hot1 | 20.14% | 7.27% | 6.12 $\times10^{-11}$ | See above for genes induced by IRAK4 |
| Sko1 | 9.17% | 25.19% | 1.34 $\times10^{-9}$ | See above for genes induced by IRAK4 |
| Adr1 | 9.60% | 19.48% | 1.97 $\times10^{-8}$ | See above for genes induced by IRAK4 |
| Ndt80 | 6.93% | 38.96% | 6.82 $\times10^{-6}$ | See above for genes induced by IRAK4 |
| **Regulating repressed genes** | | | | |
| Hap1 | 4.87% | 23.33% | 1.53 $\times10^{-7}$ | See above for genes repressed by IRAK4 |
| Fkh1 | 3.01% | 55.33% | 1.33 $\times10^{-6}$ | Regulates transcription elongation. chromatin silencing at mating loci. and gene expression in G2/M phase. |
| Spt23 | 3.34% | 40.67% | 3.5 $\times10^{-6}$ | See above for genes induced by IRAK4. |
| Fkh2 | 3.14% | 44.00% | 9.78 $\times10^{-6}$ | *FKH1* paralog. |

[a] According to the SGD (www.yeastgenome.org).

**Table S9. Oligonucleotides used in this work.**

| Name | Sequence |
|---|---|
| IRAK-4-Fw | CGGGATCCATGAACAAACCCATAACA |
| IRAK-4-Rv | GCTCTAGATTACAAAGAAGCTGTCAT |
| IRAK-1-Fw | GCTCTAGACATGGCCGGGGGGCCG |
| IRAK-1-Rv | CCCAAGCTTTTACAAGCTCTGAAATTCATC |
| IRAK-2-Fw | CGGGATCCATGGCCTGCTACATCTAC |
| IRAK-2-Rv | GCTCTAGATTACAATGTAACATCCTGG |
| Snf1-Myc Fw | TTTACATTTAACAACAAAACTAATTATGGAATTAGCCGTTAAC AGTCAAAGCAATGGATCCCATCGATTTAAAGC |
| Snf1-Myc-Rv | CGTTACGATACATAAAAAAAAGGGAACTTCCATATCATTCTT TTACGTTCCACCAGAGTGCACCAAACGACATTAC |
| Atg13-Myc-Fw | GATCAAGATGATGATCTAGTATTTTTCATGAGTGATATGAAC CTTT CTAAAGAAGGTGGATCCCATCGATTTAAAGC |
| Atg13-Myc-Rv | CACCTTTTTTTTGATTATTTTTCTTTAGTTGTGCCCTTTAAAA TAAAACTTTACCATTTGAGTGCACCAAACGACATTAC |
| Gre1-mNG Fw | GTCTGGGTCAGGAAACGATGAATATGATGATGATAGTGGG AACC AAGGCGTCTGGTCTGGTAGTTCTGGTGGTATGG |
| Gre1-mNG Rv | GCTCAAGGGTGAGACCCGCACCTCAGGCATGTAATAGAAG CTTCG ACCACCGCATACTGGATGGCGGCGTTAG |
| Gre1-C-Fw | ATGTCCAATCTATTAAACAAG |
| mNG-Rv | CCATACCACCAGAACTACC |
| IRAK-4-213-214A-Fw | AACACAACTGTGGCAGTGGCGGCGCTTGCAGCAATGGTT GAC |
| IRAK-4-213-214A-Rv | GTCAACCATTGCTGCAAGCGCCGCCACTGCCACAGTTGT GTT |
| IRAK-1-500 | CTGTGGCCTCCACCGCCATC |
| IRAK-1-1300 | CCCGGGCACCCCCGCCACC |
| IRAK-4-650 | GCAATGGTTGACATTACTAC |
| ACT1_Fw | ACGAAAGATTCAGAGCCCCA |
| ACT1_Rv | GCAGATTCCAAACCCAAAACA |
| CTT1_Fw | TCAAAAATGAAGACAACGACGAA |
| CTT1_Rv | ACAATATGCTCGTTTCTGATTTGG |
| ALD3_Fw | CACATGTTTGCTCGCGATATTAA |
| ALD3_Rv | GCTTCTTCTTGATTGGTTTGATTG |
| GRE1_Fw | GGTCGTACAAGAGGTGCTCAA |
| GRE1_Rv | CGTCTACCGCTGCCGATATT |

**Table S10. Plasmids used in this work.**

| Name | Reference |
|---|---|
| pEG(KG)-GST-Ø | This work |
| pEG(KG)-GST-IRAK4 | This work |
| pEG(KG)-GST-IRAK4(KD) | This work |

| | |
|---|---|
| pEG(KG)-GST-IRAK1 | This work |
| pEG(KG)-GST-IRAK2 | This work |
| pRS314-GFP-Atg8 | Suzuki et al., 2013 |
| pRS413-Sch9-HA | Gift of Claudio De Virgilio. University of Fribourg, Switzerland. |
| pRS423-prCUP-6xMyC-Cki12-200(S125/130A) | Deminoff et al., 2006 |
| YEplac112-Ilv6-mCherry | Fernández-Acero et al., 2019 |
| pAP67 | Gift of Adam Perez and Jeremy W. Thorner. University of California, Berkeley |

**Table S11. Antibodies used in this work.**

| Name | Dilution | Source | Catalog Number | RRID |
|---|---|---|---|---|
| Anti-GST (Z-5) | 1:1000 | Santa Cruz Biotechnology | sc459 | AB_631586 |
| Anti-GFP (JL-8) | 1:1000 | Living Colors | 632380 | AB_10013427 |
| Anti-Myc (4A6) | 1:1000 | Millipore | 05-724 | AB_11211891 |
| Anti-HA (12CA5) | 1:1000 | Sigma-Aldrich | 11583516001 | AB_514505 |
| Anti-IRAK4 | 1:1000 | Cell Signaling | 4363 | AB_2126429 |
| Anti-P-AMPKα (Thr172) | 1:1000 | Cell Signaling | 2531 | AB_330330 |
| Anti-G6PDH | 1:50,000 | Sigma-Aldrich | A9521 | AB_258454 |
| Anti-Actin (C4) | 1:1000 | MP Biomedicals | 691001 | AB_2920628 |
| IRDye 800LT Goat anti-Mouse IgG | 1:50,000 | LI-COR Biosciences | 926-32210 | AB_2687825 |
| IRDye 680LT Goat anti-Mouse IgG | 1:50,000 | LI-COR Biosciences | 926-68020 | AB_10706161 |
| IRDye 800CW Goat anti-Rabbit IgG | 1:50,000 | LI-COR Biosciences | 926-32211 | AB_621843 |
| IRDye 680CW Goat anti-Rabbit IgG | 1:50,000 | LI-COR Biosciences | 926-68021 | AB_10693472 |

### SUPPLEMENTARY MATERIALS AND METHODS

#### Cell Permeabilization and Cell Cycle Analyses by Flow Cytometry

Plasma membrane integrity was analyzed by staining with propidium iodide (PI). A 1 mL aliquot of the culture was added of PI to a final concentration of 0.0005%. The sample was gently mixed by inversion and incubated for 2 min at 30 °C in the dark. Afterwards, a 1:10 dilution was prepared in PBS, and cells were analyzed by flow cytometry.

Cell cycle analysis was performed following the protocol described by Haase & Lew (5). Approximately $10^7$ yeast cells were collected by centrifugation at 2,500 rpm for 3 min at 4 °C. For fixation and permeabilization, the pellet was resuspended in 1.5 mL PBS, followed by the addition of 3.5 mL of ice-cold molecular biology–grade absolute ethanol, reaching a final concentration of 70% (v/v). The samples were gently mixed by inversion and incubated overnight at 4 °C. Then, ethanol was removed by centrifugation, and cells were resuspended in PBS and transferred to microtubes. To eliminate RNA, cells were treated for 2 h at 37 °C with 500 µL of freshly prepared RNase solution (2 mg/mL RNase in 50 mM Tris-HCl pH 8, 15 mM NaCl), which had been boiled for 15 min and cooled prior to use. After centrifugation, cells were incubated for 15 min at 37 °C in 200 µL of a protease solution (5 mg/mL pepsin in 0.17% HCl). Finally, cells were stained with PI at a final concentration of 0.0005% in PBS, incubated in the dark at 30 °C for 2 min, diluted 1:10 in PBS, and analyzed by flow cytometry. Flow cytometry experiments were performed using a BD FACScan Scan (RRID: SCR_019596) equipped with a 488 nm excitation laser and emission filters with a bandwidth of 530/30 (FL1), 585/42 (FL2), and 670 (FL3) nm. In all cases, at least 10,000 cells per sample were analyzed. Data were processed using FlowJo (RRID: SCR_008520) software. Analyses were performed at the Flow Cytometry and Fluorescence Microscopy Unit of University of Madrid (RRID: SCR_011166).

#### Real-Time Quantitative PCR (RT-qPCR)

Gene expression levels identified by DNA microarrays were validated by real-time quantitative PCR (RT-qPCR) using the same RNA samples. The Genomics Core Facility at the Complutense University of Madrid performed cDNA synthesis and RT-qPCR (UCM; RRID: SCR_011166). For reverse transcription, 2 µg of total RNA was used with the AS Transcription System kit (Promega; RRID: SCR_006724) under the following conditions: 45 min at 42 °C, 5 min at 95 °C, and 5 min at 4 °C. The resulting cDNA was diluted 1:100 in Milli-Q water. Each 384-well reaction contained 4.5 µL of diluted cDNA, 5 µL of Power SYBR Green PCR Master Mix (Applied Biosystems; RRID: SCR_005039), and 0.6 µL of each primer (5 µM). Primer sequences are listed in **Suppl. Table S9** and were provided by J. Arroyo's lab (Department of Microbiology and Parasitology, Faculty of Pharmacy, UCM). Reactions were performed in technical duplicates using a Life Technologies QuantStudio 7 Real Time PCR System (RRID: SCR_020245) with the following cycling conditions: 10 min at 95 °C (initial denaturation), followed by

40 cycles of 15 min at 95 °C, and a final step of 1 min at 60 °C. A melting curve analysis was performed to verify the specificity of amplification. Gene expression levels were normalized to the *ACT1* housekeeping gene and to control samples carrying the empty vector pEG(KG)-GST-Ø. Relative expression changes were calculated using the 2–ΔΔCt method (6).

#### Actin Cytoskeleton Staining

Actin cytoskeleton staining was performed using rhodamine–phalloidin (Molecular Probes; RRID: SCR_013318). Cells were fixed with 8% (v/v) paraformaldehyde for 1 h at 4 °C, then centrifuged at 2,500 rpm for 3 min. They were washed three times with a freshly prepared buffer containing 0.1 M $KH_2PO_4$, 0.2 M EGTA (pH 6.9), and 0.1 M $MgCl_2$. For permeabilization, cells were resuspended in 100 µL of this buffer and 50 µL of 1% Triton X-100, mixed by inversion, and incubated for 2 min. After centrifugation at 3,000 rpm for 1 min, cells were washed three times with PBS, resuspended in 100 µL of buffer, and incubated overnight at 4 °C. Then, rhodamine–phalloidin was added to a final concentration of 2.8 ng/µL, and samples were incubated for 1 h at room temperature in the dark. Cells were then washed with PBS and analyzed by fluorescence microscopy.

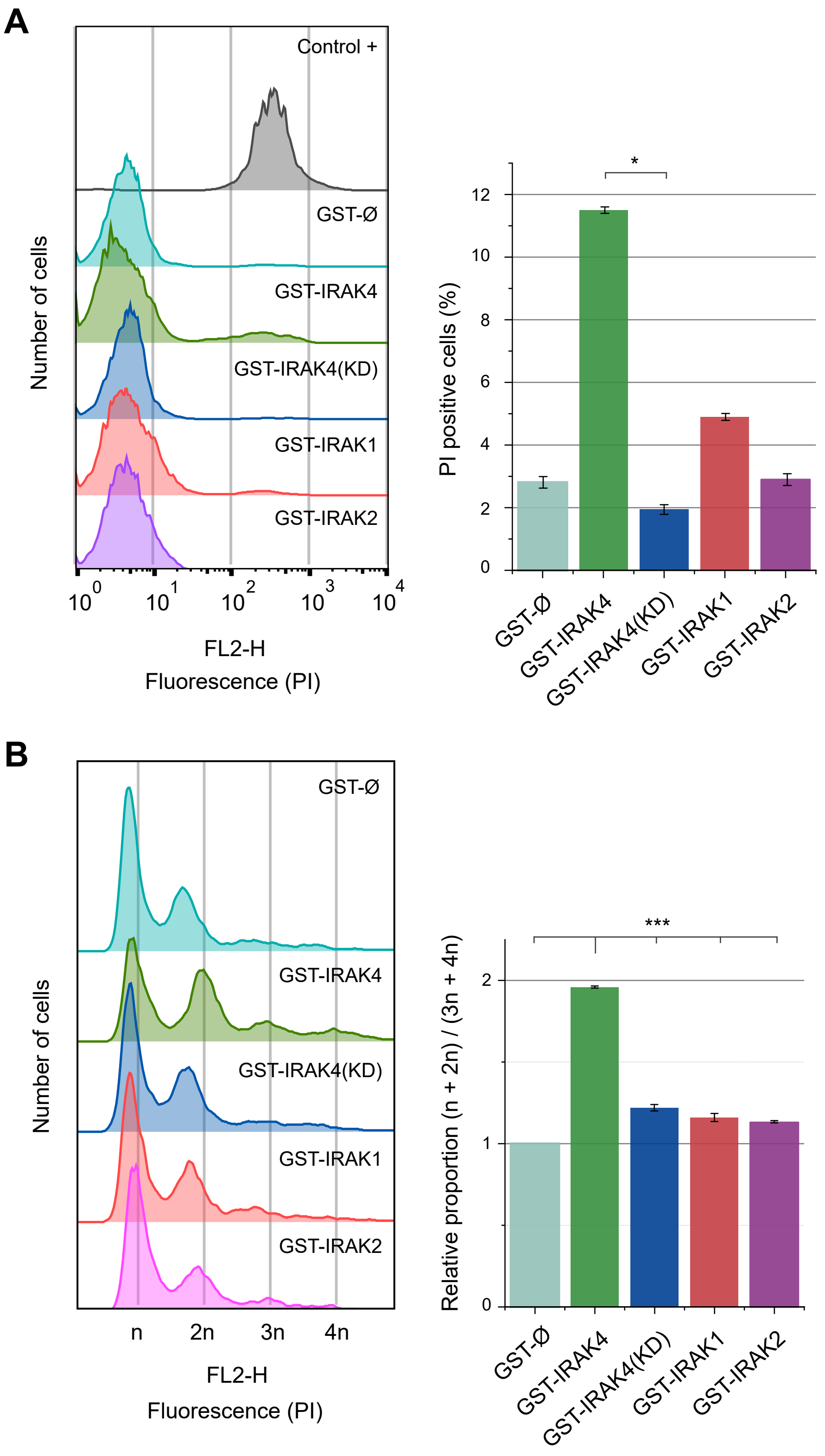


**Figure S1**

**A.** Cells of the YPH499 strain transformed with the same plasmids as in **A** were incubated for 6 h in SG Ura-Leu- medium. After PI staining, they were analyzed by flow cytometry. The left panel shows a representative histogram of each sample, and the right panel displays the corresponding bar graph of the median percentage (%) of PI-positive cells in each case. The experiment was performed with biological triplicates for each sample. Error bars represent the interquartile range Q3-Q1. The asterisk (*) corresponds to a p-value of <0.05 according to Dunn's post hoc test with Bonferroni correction.

**B.** Cells of the YPH499 strain transformed with the same plasmids as in **A** and incubated under the same conditions as in **B**. After cell fixation and PI staining, the cells were analyzed by flow cytometry. The left panel shows a representative histogram of each experimental condition. The right panel displays a bar graph representing the proportion of cells with 3n + 4n genetic content compared to n + 2n cells, normalized to the control condition with the empty plasmid (GST-Ø). The experiment was performed with biological triplicates for each sample. Error bars represent the SD. Asterisks *** correspond to a p-value of <0.001 according to Tukey's HSD test.

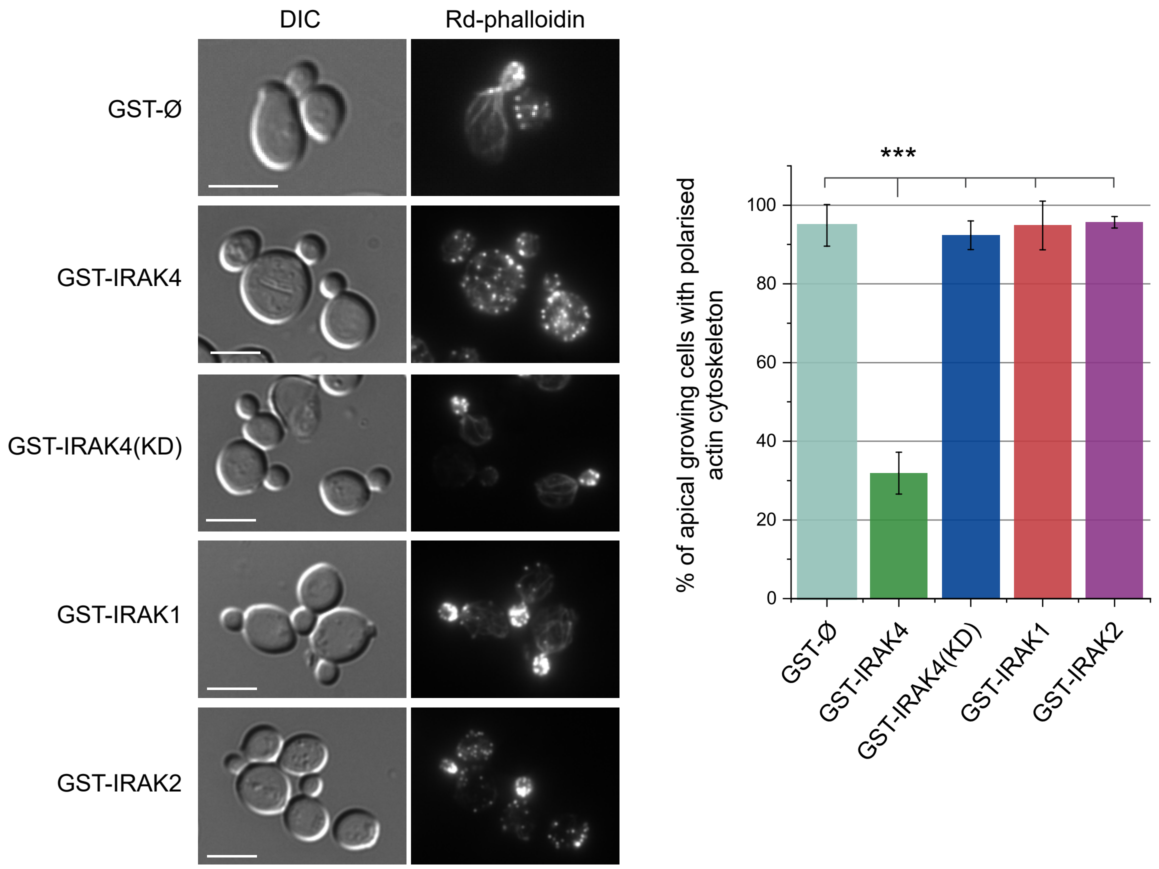


**Figure S2.**

**IRAK4 expression induces actin cytoskeleton depolarization in dividing yeast cells.**
Heterologous expression of IRAK4 in *S. cerevisiae* leads to depolarization of the actin cytoskeleton during the cell division cycle. Left panel: Differential interference contrast (DIC) and fluorescence microscopy images of YPH499 cells transformed with plasmids encoding GST alone (GST-Ø), GST-IRAK4, GST-IRAK4(KD), GST-IRAK1, or GST-IRAK2. Cells were cultured in SG Ura- Leu- medium for 8 h to induce protein expression, then fixed and stained with rhodamine-phalloidin. Images are representative of three biological replicates. Scale bar: 5 µm. Right panel: Quantification of apically growing cells displaying polarized actin cytoskeleton. Over 150 cells were counted per condition. Bars represent the mean percentage ± standard deviation (SD). Statistical significance is indicated as *** $p < 0.001$ (Tukey's HSD test).

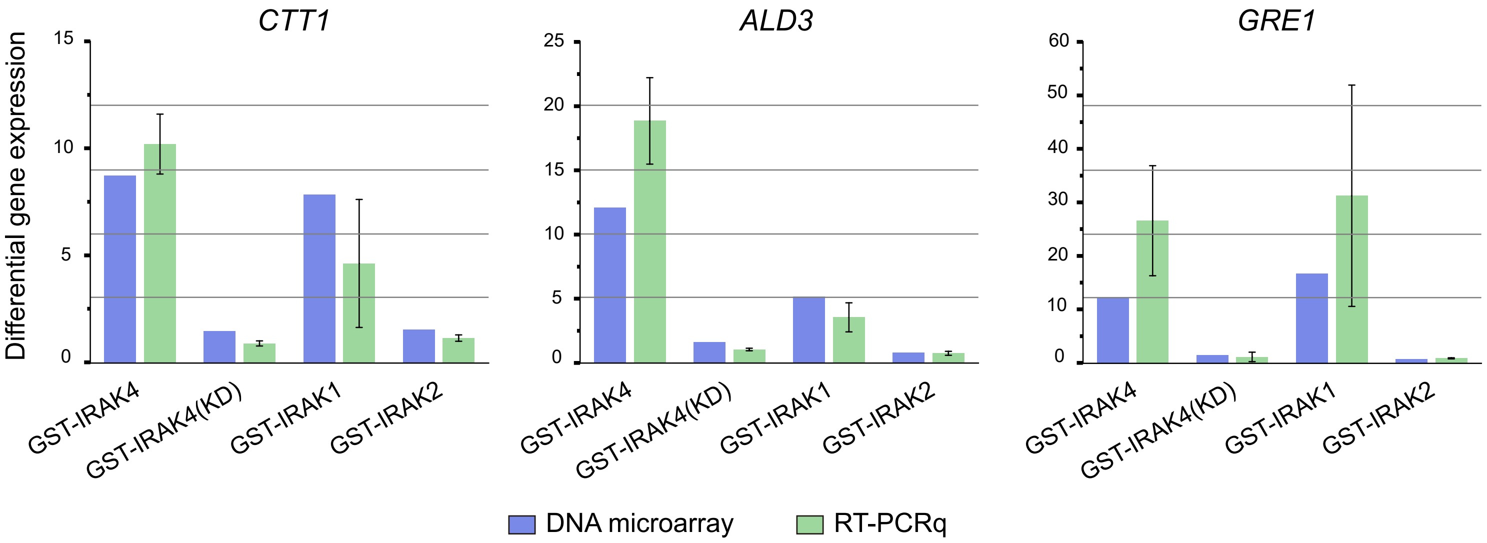


**Figure S3**

Validation of transcriptomic data obtained from DNA microarrays by qRT-PCR. Bar graphs showing in blue the differential gene expression ratios, or fold change (FC), of each gene (CTT1, ALD3 or GRE1) for each experimental condition [IRAK4, IRAK4(KD), IRAK1 and IRAK2] compared to the control condition (Ø) obtained from DNA microarrays. In green, the gene expression level of each gene for each experimental condition normalized with respect to the constitutively expressed gene ACT1 and the control condition, obtained by qRT-PCR and the 2-ΔΔCt method described by Livak & Schmittgen, 2001. In the case of qRT-PCR, the results shown are the arithmetic mean of the biological triplicates, except for the GRE1 gene in IRAK1, where one biological duplicate was used. RT-PCRq was performed in duplicate for all clones. Error bars represent SD.

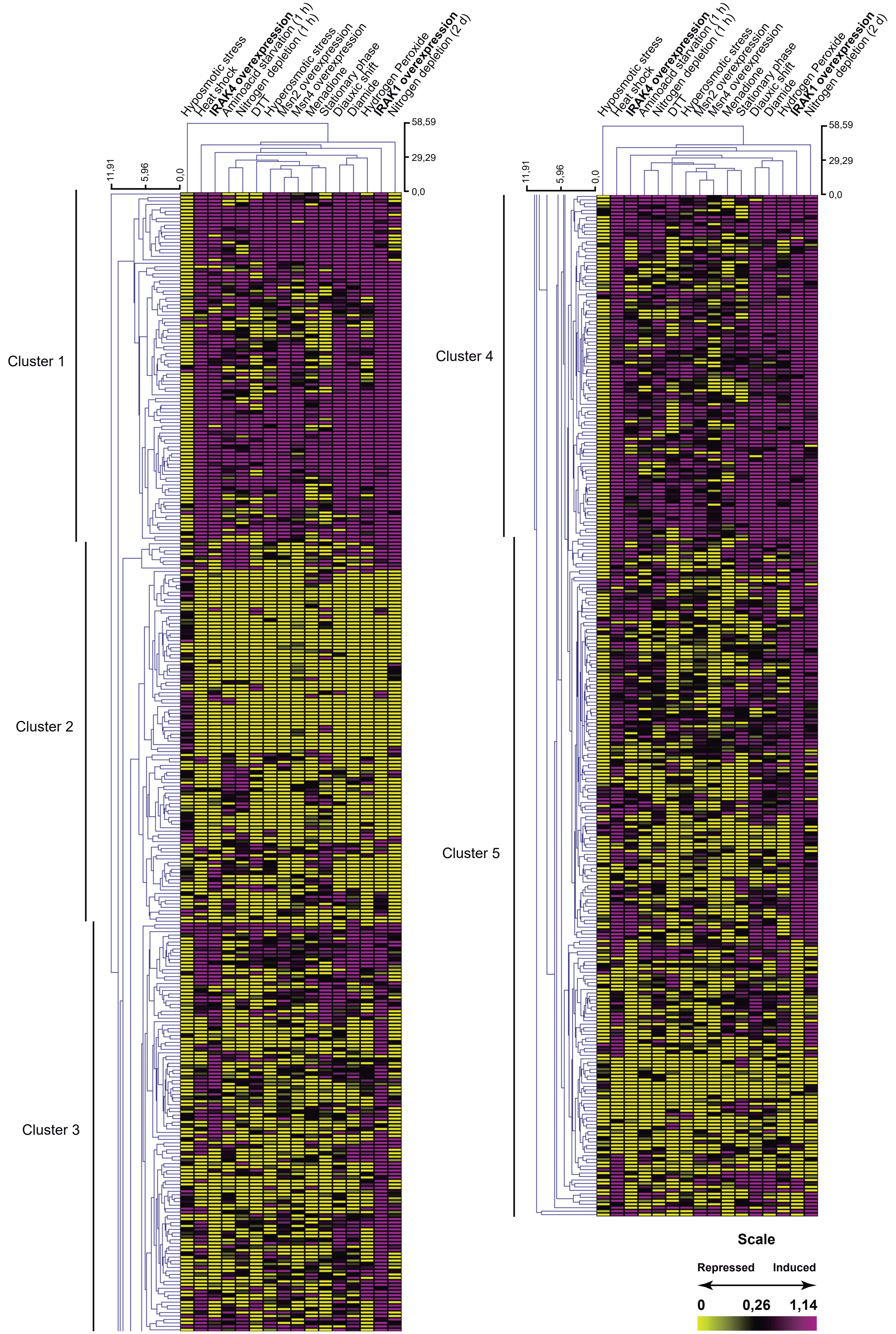

Hyposmotic stress
Heat shock
IRAK4 overexpression
Aminoacid starvation (1 h)
Nitrogen depletion (1 h)
DTT
Hyperosmotic stress
Msn2 overexpression
Msn4 overexpression
Menadione
Stationary phase
Diauxic shift
Diamide
Hydrogen Peroxide
IRAK1 overexpression
Nitrogen depletion (2 d)
58,59
29,29
0,0
11,91
5,96
0,0
Cluster 1
Cluster 2
Cluster 3
Cluster 4
Cluster 5
Scale
Repressed
Induced
0
0,26
1,14

**Figure S4**

**Clustering of *S. cerevisiae* transcriptional profiles under various stress conditions.**
Gasch et al. (2000) analyzed the yeast transcriptional response to a variety of stress conditions at different time points. For this clustering analysis, the following conditions were selected: heat shock (30 min), hypoosmotic stress (30 min), amino acid starvation (1 h), nitrogen depletion (1 h), nitrogen depletion 2 days), diauxic shift (18.5 h), entry into stationary phase in YPD (8 h), overexpression of MSN2, overexpression of MSN4, and exposure to: hydrogen peroxide ($H_2O_2$, 0.32 mM, 30 min), menadione (1 mM, 120 min), DTT (2.5 mM, 120 min), diamide (1.5 mM, 30 min), and sorbitol (1 M, 30 min; hyperosmotic stress). For each condition, the time point capturing the peak transcriptional response was selected. In cases with a clear temporal shift in expression, two representative time points were included. For IRAK4 and IRAK1, the analysis considered the set of differentially expressed genes shared between both conditions. The graphical representation follows a tabular heatmap format, where each row represents a gene and each column corresponds to a specific experimental condition. Two dendrograms display the relationships between genes (vertical axis) and between conditions (horizontal axis), with branch lengths indicating the degree of similarity in expression profiles. Transcript abundance variations are visualized using a color scale, where purple and yellow tones indicate, respectively, higher and lower expression ratios compared to the corresponding control. The saturation level of the color reflects the magnitude of the expression ratio. Expression ratios were $\log_2$-transformed for this analysis.

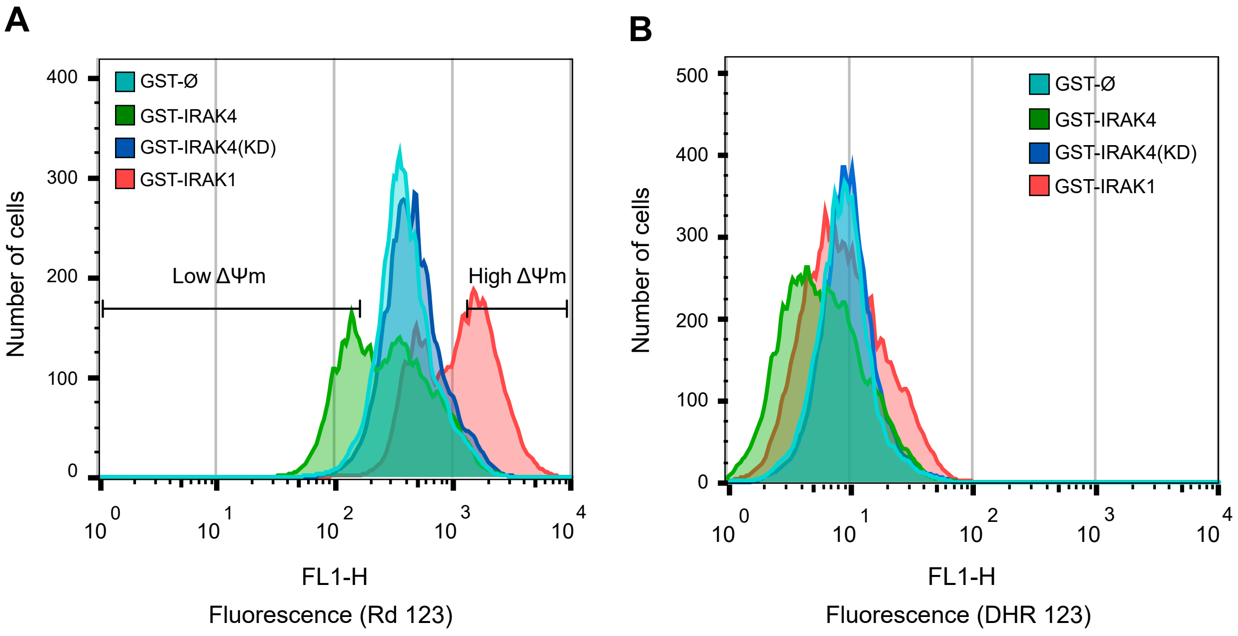


**Figure S5.**

**A.** Superimposed histogram of the fluorescence signal emitted by Rd 123. Cells of strain YPH499 were transformed with the plasmids pEG(KG)-GST-Ø, pEG(KG)-GST-IRAK4, pEG(KG)-GST-IRAK4(KD), or pEG(KG)-GST-IRAK1. Expression of the heterologous proteins was carried out for 12 h in SG Ura-Leu- culture medium. The experiment was performed in biological triplicates, and a representative result is shown. 10,000 cells from each transformant were analyzed. The intensity of fluorescence emitted by Rd 123 is shown on the abscissa axis, and the number of cellular events on the ordinate axis. The fluorescence signal from dead cells was eliminated from the analysis by co-staining with PI (propidium iodide). **B.** Overlaid histogram of the fluorescence signal emitted by DHR 123. Cells of strain YPH499 were transformed with the plasmids pEG(KG)-GST-Ø, pEG(KG)-GST-IRAK4, pEG(KG)-GST-IRAK4(KD), or pEG(KG)-GST-IRAK1. Expression of heterologous proteins was carried out for 12 h in SG Ura-Leu- medium. The experiment was performed with biological triplicates, and a representative result is shown. 10,000 cells from each clone were analyzed. The fluorescence signal from dead cells was eliminated from the analysis by co-staining with PI.
